# A Precision Gene Engineered B Cell Medicine Producing Sustained Levels of Active Factor IX for Hemophilia B Therapy

**DOI:** 10.1101/2025.04.06.647090

**Authors:** Hanlan Liu, Swati Singh, Timothy J Mullen, Caroline Bullock, Sean Keegan, Troy Patterson, Sakshisingh Thakur, Amy Lundberg, Sol Shenker, Ryan Couto, Charuta Yadav, Shamael Dastagir, Lily Li, Wayne Bainter, Ella Liberzon, Connor R Malloy, Cicera R Lazzarotto, Toshiro K. Ohsumi, Shalini Chilakala, Huei-Mei Chen, Rashmi Kshirsagar, Anja F Hohmann, Sean Arlauckas, Adam Lazorchak, Chris Scull, Richard A Morgan

**Author notes:** Corresponding Author Name: Hanlan Liu, Address: One Kendall Square, B200, 3^rd^ Floor, Cambridge, MA 02139, USA.

## Abstract

Hemophilia B gene therapy treatments currently have not addressed the need for predictable, durable, active, and redosable factor IX (FIX). Unlike conventional gene therapy, engineered B Cell Medicines (BCMs) are durable, redosable, and titratable, and thus have the potential to address significant unmet needs in the Hemophilia B treatment paradigm. BE-101 is an autologous BCM comprised of expanded and differentiated B lymphocyte lineage cells genetically engineered *ex vivo* to secrete FIX-Padua. CRISPR/Cas9 mediated gene editing at the C-C chemokine receptor type 5 locus was used to facilitate transgene insertion of an AAV6-encoded DNA template via homology-directed repair. Transgene insertion did not alter B cell biology, viability, or differentiation into plasma cells. Appreciable levels of BE-101-derived FIX-Padua were detected within 1 day after IV administration in mouse and steady state was reached within 2 weeks and persisted for over 184 days. Redosing produced an increase in FIX-Padua production close to linear dose proportionality. Comprehensive genotoxicity analysis found no off-target issues of concern. No safety signals were observed in animal tolerability and GLP toxicology studies. In conclusion, BE-101 produces sustained levels of active FIX-Padua with the ability to engraft without host preconditioning and with the potential for redosing and titratability.

## Introduction

Mutations in the *F9* gene coding for factor IX (FIX) lead to FIX deficiency and severe bleeding events in people with hemophilia B, (PwHB). Recommended therapy is prophylactic infusion of exogenous FIX,^1,2^ however, the short biological half-life requires frequent administration once weekly to biweekly to maintain therapeutic levels. Even extended half-life FIX prophylaxis may require individual pharmacokinetic monitoring for optimal outcomes.^3^ Two recently approved hemostatic rebalancing agents, marstacimab and fitusiran, do not replace the missing FIX, rather reduce Tissue Factor Pathway Inhibitor (TFPI) and antithrombin, respectively, still require weekly or bimonthly subcutaneous administration.^4,5^ A considerable barrier to treatment adherence for PwHB that leads to breakthrough bleeding events is the life-long need for frequent infusions at intervals in which trough levels are low.^6,7^ Real-world outcome studies demonstrate that both standard and extended half-life FIX prophylaxis are associated with breakthrough bleeding in PwHB (87/108 (81%) and 23/34 (68%) individuals, respectively, reported bleeding events in a 12 mo period).^8^

Adeno-associated virus (AAV) vector-based gene therapy products (etranacogene dezaparvovec and fidanacogene elaparvovec-dzkt) are approved for adults with HB and show superiority versus FIX prophylaxis with respect to annualized bleed rates.^9,10^ While promising, these therapies carry potential risks of systemic and liver toxicity as well as genotoxicity due to random AAV chromosomal insertions. Furthermore, AAV immunogenicity precludes re-dosing and may decrease therapeutic efficacy.^11,12^ Significantly, AAV-based *in vivo* gene therapy is not approved for children due to potential waning of sufficient FIX activity with hepatic turnover and liver growth as well as unknown safety risks. Thus, significant unmet medical needs remain for new therapies providing predictable, durable, and redosable FIX, especially in children where early intervention is critical to prevent joint damage.^1^

An autologous source of stable and durable FIX is a promising therapeutic goal to address unmet needs for PwHB. Antibody producing long-lived plasma cells (PCs) are attractive *ex vivo* cell therapies. Derived from antigen-driven differentiation of B cells, PCs are inherently long-lived (half-lives ranging from 6 months to decades)^13,14^ and actively home to and reside in the bone marrow compartment where they secrete high levels of antibodies (>1,000 antibodies/cell/second).^15^ Terminally differentiated human PCs derived from genetically engineered B cells (B Cell Medicines, BCMs) produce sustained and high levels of transgenic proteins *in vivo* in immunodeficient mice without pre-conditioning.^16^ In non-human primates, PET/CT tracking of zirconium-89-oxine labeled, *ex vivo* differentiated plasma cells show that the cells rapidly home to, and engraft in, the bone marrow, spleen and liver after infusion into unconditioned autologous subjects.^17^ Moreover, in a re-dosing experiment using this approach, the biodistribution was identical to what was observed following the initial dose demonstrating the feasibility of redosing *ex vivo* differentiated plasma cells.^17^ CRISPR-Cas9 genome editing enables BCMs to secrete therapeutic proteins with long-term *in vivo* engraftment in humanized mouse models. In contrast to gene therapy, BCMs have the potential for redosing, making them an attractive platform for the sustained supply of biologics where continuous and adaptive dosing is required to achieve therapeutic benefit, for example, in children with hemophilia B.

In this paper we describe pre-clinical studies characterizing an autologous *ex vivo* precision gene edited cell therapy (termed BE-101) comprised of expanded and differentiated B lymphocyte lineage cells that have been genetically engineered to express and secrete FIX-Padua.

## Results

### *F9* transgene integration does not alter BCM biology, viability, or differentiation

BCM phenotype at the end of the culture process (**Figure 1A**) was compared for BE-101 cells and BCMs electroporated with RNP without subsequent AAV transduction. Engineering was confirmed by *CCR5* locus editing (∼80%, **Figure 1B**) and *F9* transgene integration (∼15-20%, **Figure 1C**). FIX-Padua production was evident by day 6 and remained through day 13 (**Figure 1D**). Neither viability nor phenotype differed between BE-101 cells and control cells electroporated with RNP alone. (**Figure 1E-1H**).

**Figure 1:**
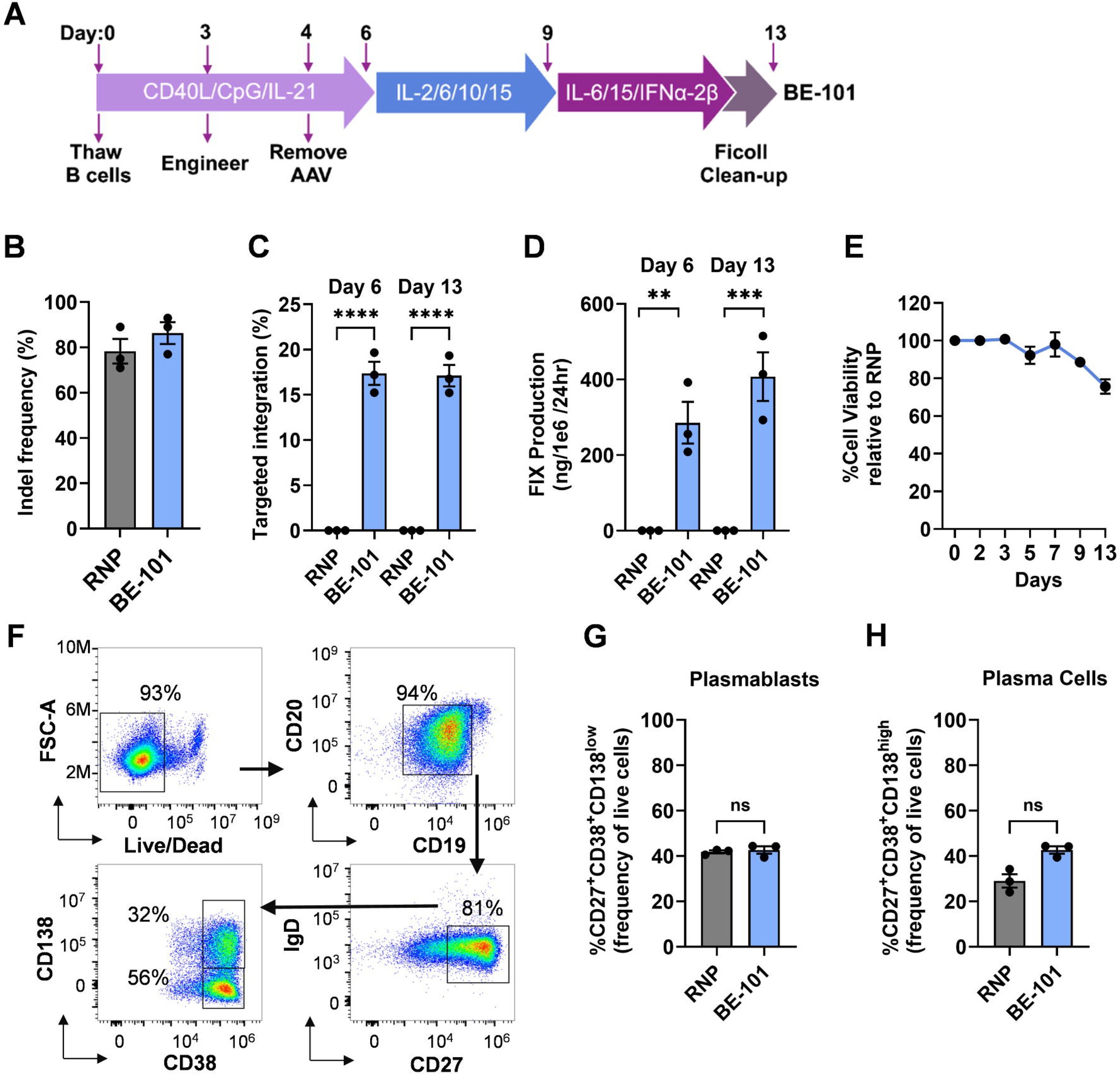
Impact of engineering on BCM viability, phenotype, and FIX-Padua production. (A) Diagrammatic representation of B cell culture and differentiation process in vitro. (B) Percentage of cutting frequency at the CCR5 locus, measured by Inference of CRISPR Edits (ICE), of RNP-only or BE-101 engineered BCM. (C) Percentage of transgene-containing alleles at days 6 and 13, determined by ddPCR, comparing BE-101 to the RNP-only control. (D) FIX-Padua production (ng/10^6^ cells/24 hr), determined by ELISA, at days 6 and 13. (E) % Cell viability during the B cell differentiation process relative to RNP-only control. (F) Representative flow cytometry data from day 13 BCMs, illustrating the gating strategy. (G) and (H) Percentage of plasmablasts (CD27+ CD38+CD138^low^) and plasma cells (CD27+CD38+CD138^high^) among the total live cells. The data is a repetitive experiment for n=3 donors. Error bars represent mean ± standard deviation p**** <0.0001, p***<0.0002; p**< 0.001, p*< 0.01 by one-way ANOVA; Holm-Sidak correction for multiple comparisons

### Production of functional FIX-Padua protein by BE-101 *in vitro*

BE-101 products generated from seven B cell donors secreted an average of 634 ng/10^6^ cells/day of FIX-Padua protein, approaching 45% of concomitant plasma cell IgG secretion rate at a molar level (**Table 1**). The measured FIX-Padua production rate *in vitro* was estimated at ∼540 molecules/cell/second.

**Table 1.** *In vitro* secretion rate of FIX-Padua protein in timed culture experiment in comparison with human IgG (hIgG) from BE-101.

| **Donor #** | **hIgG secretion rate**<br>**(ng/10<sup>6</sup> cells/24h)** | **FIX-Padua secretion rate**<br>**(ng/10<sup>6</sup> cells/24h)** | **Secretion rate of**<br>**FIX-Padua vs. hIgG**<br>**(%)<sup>a</sup>** |
| --- | --- | --- | --- |
| D1 | 2690 | 515 | 50% |
| D2 | 2266 | 293 | 34% |
| D3 | 5220 | 414 | 21% |
| D4 | 2954 | 802 | 71% |
| D5 | 3798 | 696 | 48% |
| D6 | 5124 | 855 | 44% |
| D7 | 5185 | 867 | 44% |
| **Mean (SD)** | **3891 (1287)** | **634 (229)** | **45% (15%)** |

<sup>a</sup>: calculated as FIX-Padua secretion rate divided by hIgG secretion rate and normalized by MW difference of human IgG (i.e., 150 kDa) and human FIX-Padua protein (i.e., 57 kDa).

BE-101-derived FIX-Padua protein function *in vitro* was determined in *ex vivo* cell cultures in the presence of vitamin K.^18^ Peripheral blood B cells isolated from three donors were engineered with menadione sodium bisulfite (synthetic vitamin K3). The mean amount of FIX-Padua protein produced was 407.3±111.4 ng/mL (**Figure 2A**). Vitamin K-dependent active FIX-Padua protein in the range of 1.82 to 11.6 mIU/mL corresponded to an activity/mg of protein of 15.8 to 25.4 IU/mg (**Figure 2B**). In the aPTT assay, purified protein cultured in the presence of vitamin K altered clotting time in a concentration dependent manner with % FIX-Padua activity at 28.9%, 15.4%, and 26.4% for donor A, B, and C, respectively, using 5 μg of BE-101 purified protein (**Figure 2C-2D**). Activity levels observed in FIX-Padua deficient plasma in the aPTT assay corresponded to a specific activity of 26.7 to 65.2 IU/mg of protein. Minimal activity in the absence of vitamin K was observed. Finally, LC-MS/MS analysis of trypsin-digested FIX protein derived from BE-101 cultured in the presence of vitamin K for 48 hours showed up to 11 out of 12 possible glutamic acid residues in GLA domain underwent γ-carboxylation (**Figure 2E**).

**Figure 2.**
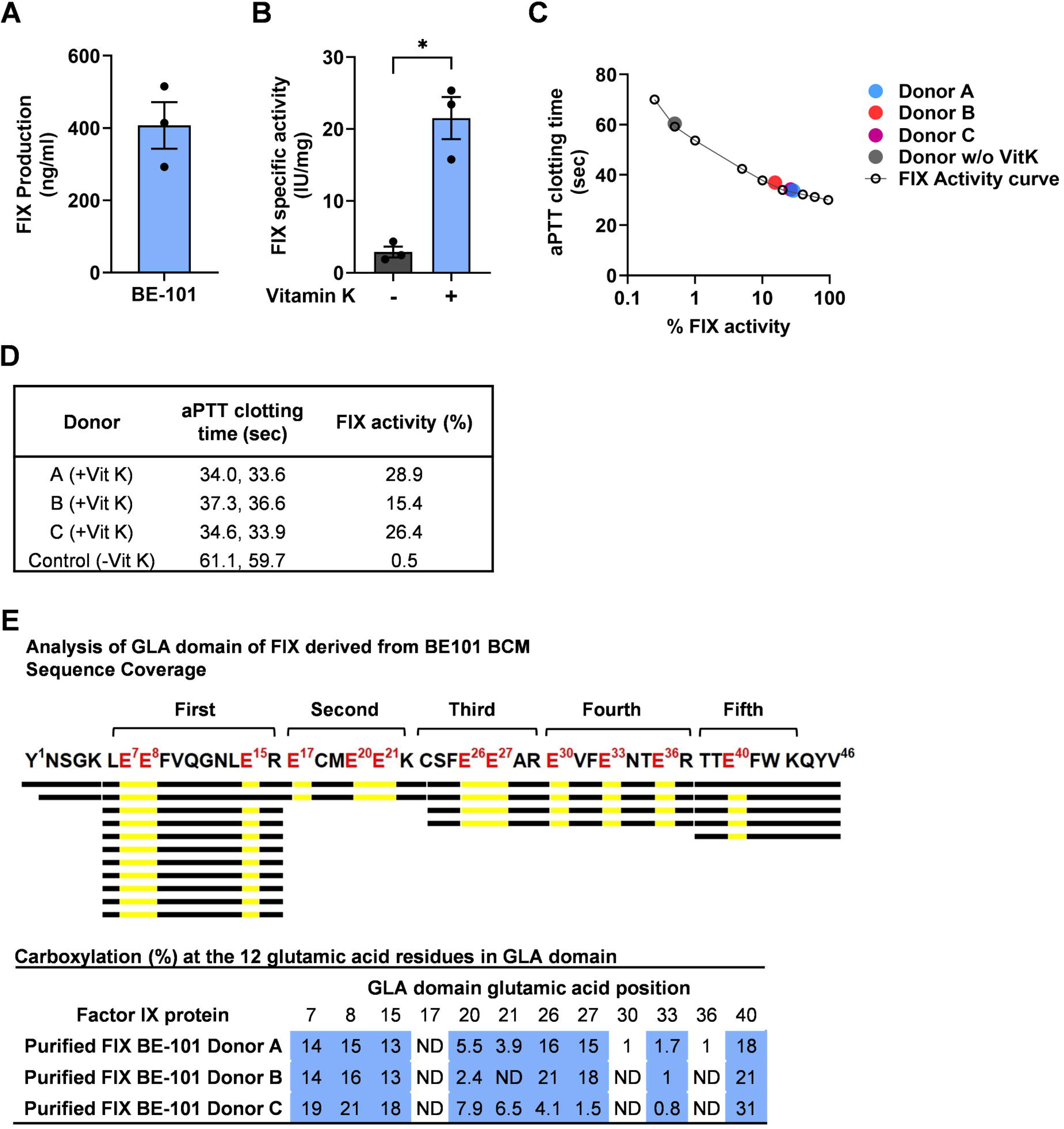
*In vitro* FIX-Padua protein secretion and activity from BE-101. **(A)** FIX-Padua protein levels in supernatant production from engineered B cells on day 13 of culture as measured by ELISA. The bar graph depicts mean and standard deviation of 3 donors **(B)** Active FIX-Padua protein production by BE-101 culture in the presence of vitamin K, added for 48 hours from day 11-13. Activity was measured from culture supernatant using a human FIX selective antibody capture step coupled with activity determination using Rossix FIX activity assay. The bar graph represents the calculated mean specific activity and standard deviation of 3 donors. **(C)** FIX-Padua activity was measured using activated partial thromboplastin time using PTT-automate activation and measurement of clotting times on the Stago Satellite. FIX-Padua activity curve was made from dilutions of normal pooled plasma into hemophilia B plasma. FIX-Padua protein purified from engineered B cell cultures at 5 μg/mL, was measured for Donors A to C (+ Vit K, in the presence of vitamin K) and one control donor (-Vit K, in the absence of Vitamin K). **(D)** aPTT times from panel C, Donors A to C. **(E)** LC MS/MS peptide coverage of FIX GLA domain after trypsin digestion with corresponding glutamate carboxylation status.

### Engineering efficiency, FIX-Padua protein production, and cell phenotype of BE-101 produced from the scaled-up process

After demonstrating *F9* transgene integration does not alter BCM biology, viability, or differentiation and production of functional FIX-Padua protein by BE-101 *in vitro*, we optimized gene editing and *ex vivo* culture methods using the scaled-up process. Parameters varied included the electroporation conditions and timing, sgRNA to Cas9 ratio, RNP concentration, and cell density during electroporation. Optimization resulted in an average of 88% cutting frequency at *CCR5* locus in activated primary human B cells from 10 donors (**Figure 3A**). An average 51% *F9* gene integration was measured by ddPCR after optimizations were made (**Figure 3B**), which represented a ∼2.5-3-fold boost in engineering efficiency (**Figure 1C**) with a concomitant average increase in FIX production of 1343.4 ng/mL collected (**Figure 3C**). These optimized procedures reproducibility yielded 95% CD38+ BCMs from five independent donors comprised of approximately 20% CD27+/CD38+/CD138^high^ plasma cells and approximately 60% CD27+/CD38+/CD138^low^ measured from 4 out of the 5 independent donors (**Figure 3D and 3E, Table S1 and S2**).

**Figure 3:**
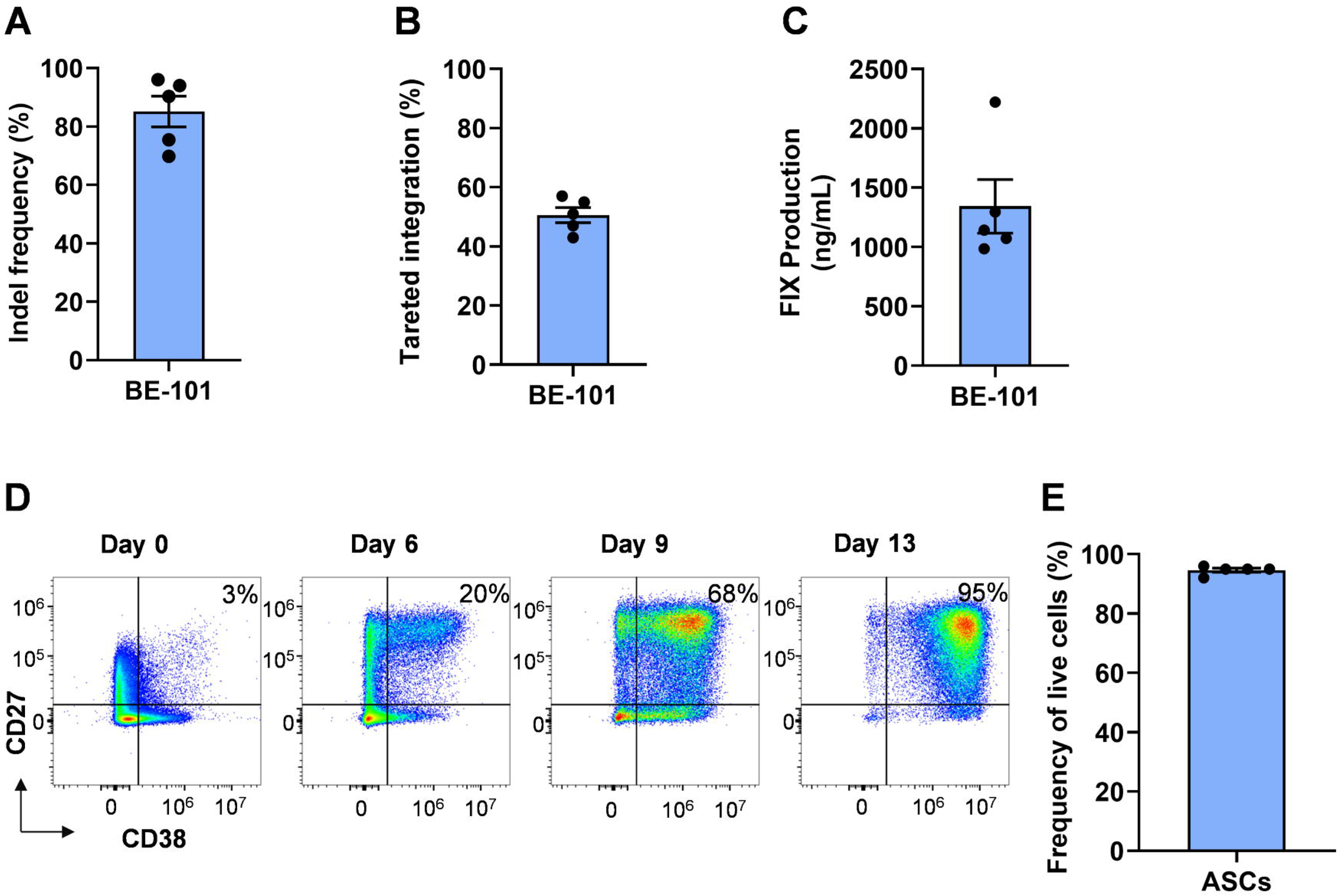
Engineering efficiency, FIX-Padua production, and cell phenotype at the scale-up process. (A) Percent cutting frequency at the *CCR5* locus. (B) Percent targeted integration (HDR) of the *F9* transgene at the *CCR5* locus. (C) Engineered B cell Factor IX (FIX) production during the final stage of cell culture, as measured in the harvest supernatant using a FIX ELISA. (D) Representative flow cytometry plot illustrating the differentiation of B cells at various time points during the B cell differentiation process. (E) Percentage of antibody secreting cells (CD38+) among live cells during and at the end of the culture process (day 13). Error bars represent the standard deviation (±) for 5 donors and 5 individual cell expansion experiments.

### Analysis of on- and off-target editing and overall genome integrity in BE-101 cell products

Following optimization of the gene editing reaction for transgene insertion efficiency at the *CCR5* on-target site, we characterized the genome integrity in BE-101 cells. First, we used both computational and experimental methods to comprehensively nominate candidate off-target sites for the gCCR5_232 guide RNA. The CRISPRitz algorithm^19^ in association with a CFD score threshold of ≥0.2^20^ identified 422 NGG-PAM containing human genomic sites of homology to the gCCR5_232 on-target sequence (with no more than four nucleotide mismatches and two bulges). Additionally, we performed G-GUIDE (a modified version of GUIDE-Seq^21^) and SITE-seq,^22^ to detect whether Cas9, complexed with GMP-grade gCCR5_232 sgRNA, cleaved off-target sites in primary human B cells and in vitro, respectively. G-GUIDE nominated 9 candidate off-target sites that were found reproducibly across technical duplicates in at least one out of 5 tested B cell donors, and as expected in primary human B cells, double strand breaks were detected in the switch regions of the IgH locus as well. SITE-seq nominated 120 candidate off-target sites that were found in at least two out of six B cell donors tested (with no more than 7 nucleotide mismatches and 5 bulges relative to the gCCR5_232 on-target site). Interestingly, the 422 sites nominated in silico and the sites nominated experimentally shared only 22 sites in common, with three sites identified by all three methods, one of which was the intended on-target site of gCCR5_232 (**Figure 4A**). Off-target sites identified by the three discovery efforts are listed in **Table S3**.

**Figure 4:**
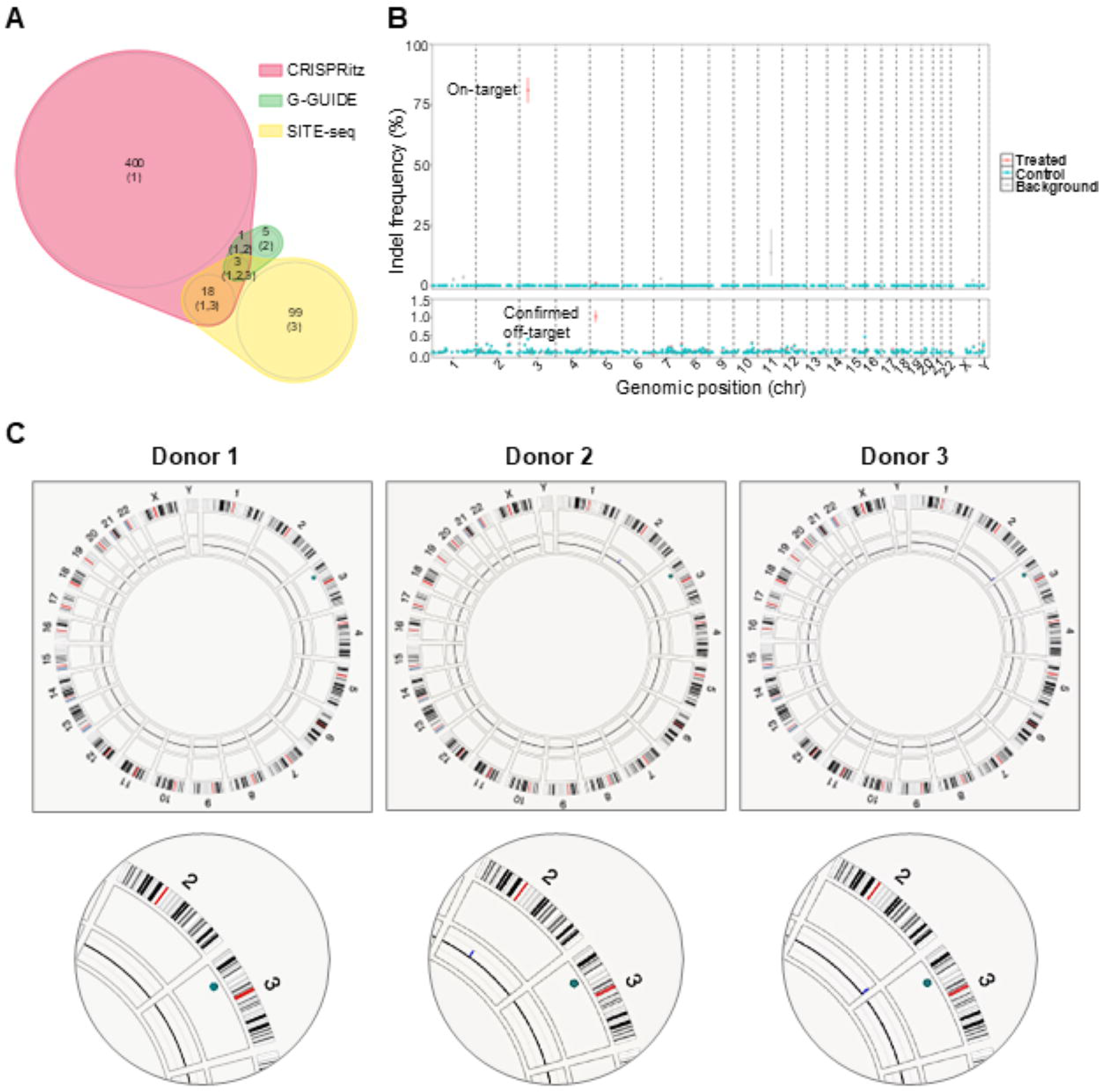
Analysis of on- and off-target editing and overall genome integrity in BE-101 cell products. (A) Venn diagram showing the number of potential gCCR5_232 off-target sites discovered by CRISPRitz, G-GUIDE, and SITE-Seq. GMP-grade gCCR5_232 was used for the nomination of sites via G-GUIDE and SITE-Seq. (B) Mean DNA editing rates +/- SEM at 457 nominated sites across the genome as measured via targeted amplicon sequencing in five lots of BE-101 cells (Treated) and unengineered, donor-matched controls (Control) plotted on a scale between 0-100% (top graph) or 0-1.5% (bottom graph). The x-axis represents chromosomal location. Sites with editing rates >1% in control samples are marked in grey (Background, BG). (C) Circus plots depicting structural variants (SVs) as identified by Bionano Rare Variant Analysis (RVA) in three lots of BE-101 (top) with magnification of the on-target insertion site on chromosome 3 (below). The outermost numerical track corresponds to the chromosome number. The cytoband information is shown in the black-and-white banding pattern with red lines indicating the centromere. The next circle indicates SVs that were uniquely identified in engineered cells but not donor-matched unengineered controls. Insertion events are represented by green dots. The next circle shows copy number variations. The inner space indicates translocation events found uniquely in engineered cells.

To determine bona fide off-target sites in BE-101 cells, DNA editing rates at the nominated loci were compared in five batches of BE-101 cells and donor-matched unengineered controls. Of the 526 candidate off-target sites, 457 were amenable to targeted DNA sequencing and reached the required minimal sequencing depth of 5000 reads per site to allow for detection of differential editing frequencies as low as 0.2%. We observed high on-target editing at *CCR5* (80.58% INDELs, p=0), and off-target activity was validated at only one candidate off-target site (0.97% INDELs, p=2.18E-46) (**Figure 4B**), located in an intergenic region on chr5.

To assess genome integrity in BE-101 cells, we performed genome-wide unbiased Optical Genome Mapping (OGM) in three lots of BE-101 cells and donor-matched unengineered controls. *De Novo* Assembly of data obtained by OGM identified insertion events of the size expected for single copy *F9* transgene integrations at the *CCR5* on-target locus with frequencies of ∼45%, in line with ddPCR results in these cells (**Figure S1, Table S4**) confirming HDR-mediated insertion of the full length FIX-expression cassette. Rare variant analysis (RVA) of the same data identified only one type of SV that was uniquely found in BE-101 cells but not donor-matched unengineered controls: large insertions at the on-target locus (detected with low molecule read counts, 5 to 25) in all three lots of BE-101 (**Figure 4C, Table S5**). The sizes of these large insertions increased in a stepwise fashion consistent with potential concatemeric AAV insertions (**Table S5**). No other reproducible SVs were identified that reached the required minimum of five molecules to support an SV identification.

### Rapid homing of BE-101 to bone marrow with stable engraftment and FIX-Padua activity in the plasma of NOG-hIL6 mice

To study the biology of BE-101 in a small animal model we used NOG-hIL6 mice. IL-6 is a pleiotropic cytokine primarily produced by antigen presenting cells and is also known to drive ASC differentiation and survival in multiple models.^23^ Murine IL-6 does not efficiently activate the human IL-6 receptor (IL-6R),^24^ thus, hIL-6 *in vivo* is required for the efficient engraftment of human BCMs.^23^ Biodistribution of BE-101 was assessed using a qPCR assay designed to detect and quantify the human-specific *Alu* sequence as a surrogate for human genomic DNA in mouse organs, bone marrow cells, and whole blood samples following BE-101 dosing in NOG-hIL6 mice. After a single IV administration of BE-101 at 20×10^6^ viable cells/mouse, human DNA was detected in femur and spinal column of NOG-hIL6 mice as early as day 1 and remained at comparable levels on day 28 (**Figure 5A and 5B**), consistent with the characteristics of plasma cells homing to the bone marrow niche. An equivalent number of human cells persisted in femur and spinal column on day 28 was estimated to be less 1% of cells dosed using 6.41 pg DNA per human cell from a male donor.^25^ Human DNA levels decreased significantly or were found to be below the limit of quantitation by day 28 in all non-bone marrow compartment tissues examined including whole blood, spleen, kidney, lung, liver, heart, brain, and testis (**Figure 5C**). Similar homing and engraftment were observed using surrogate luciferase-engineered BCMs, determined by *in vivo* bioluminescent imaging (BLI) (**Figure 5D-5F**). Among all non-homing tissues evaluated, apparent clearance of human DNA in kidney was not as rapid as other tissues. However, the *Alu* qPCR assay cannot discriminate fragments of human DNA from DNA isolated from intact human cells. It is known that the main organ for clearance of oligonucleotides^26^ is the kidney and thus it is possible that the *Alu* specific signal detected in kidney at later time points originated from fragments of human DNA.

**Figure 5.**
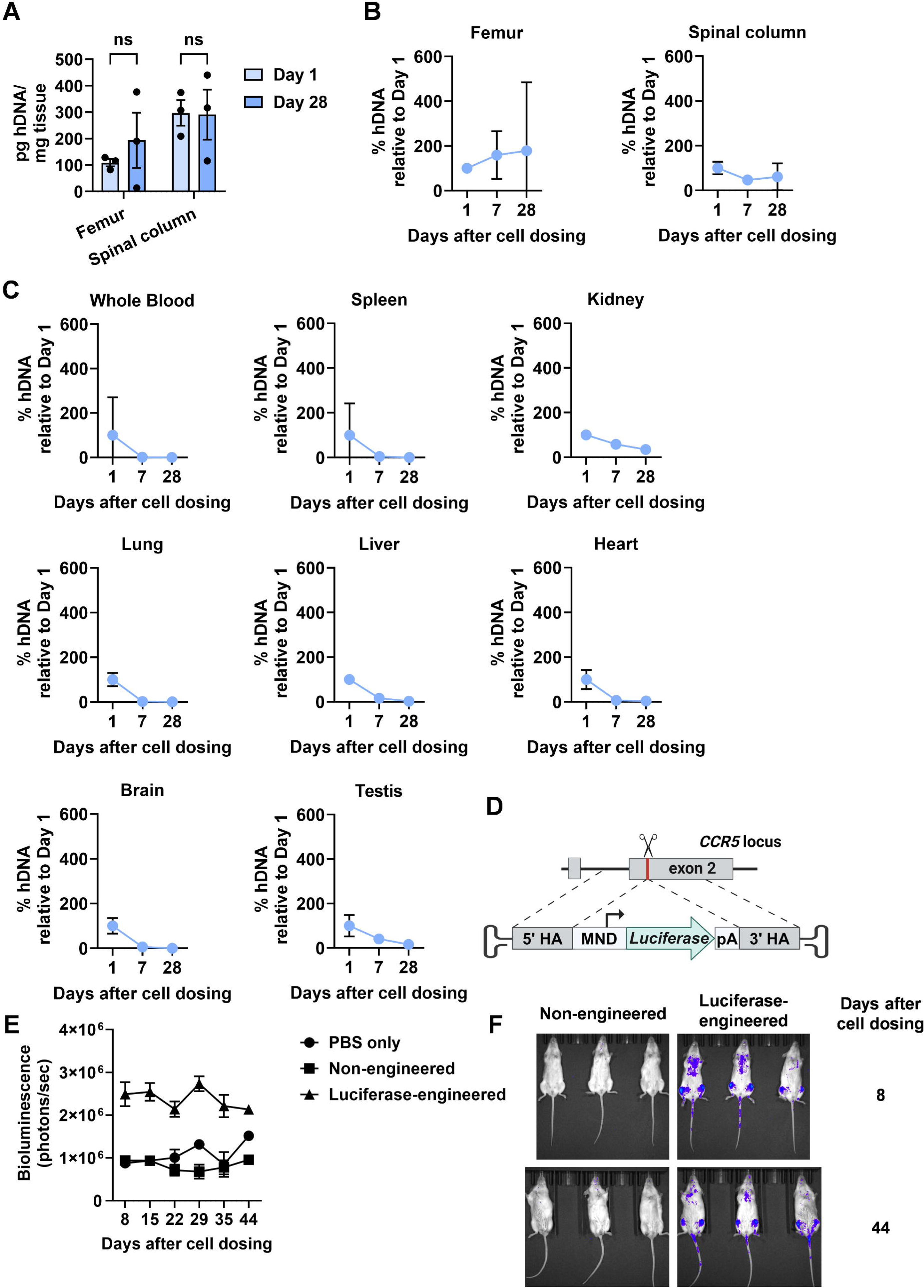
Bone marrow trafficking and stable engraftment of BE-101 in NOG-hIL6 Mice. Biodistribution of BE-101 in NOG-hIL6 mice was measured by a quantitative polymerase chain reaction (qPCR) method assay designed to detect and quantify the human-specific *Alu* sequence as a surrogate for human genomic DNA in mouse organs, bone marrow cells, and whole blood samples. (A) Mean human DNA mass detected in femur and spinal column of NOG-hIL6 mouse mice on day 1 and day 28 post IV administration of BE-101 at 20 x 10^6^ viable cells/mice. Error bars represent ± standard error of the mean for n = 3 mice on day 1 and day 28. Mann-Whitney U test was used to assess statistical significance; ns = not significant. (B) and (C) Shown are the mean percentages of human genomic DNA detected in (B) bone marrow-containing tissues and (C) whole blood and various NOG-hIL6 mouse organs on day 7 and day 28 relative to the day 1 post IV administration of BE-101 at 20 x 10^6^ viable cells/mice. Error bars represent ± coefficient of variation for n = 3 mice at each time point. (D) Schematic of the firefly luciferase transgene for targeted integration at the *CCR5* locus. (E) Quantification of bioluminescence signal over time. Each data point represents the mean and the error bars represent ± SEM for n = 3 mice in each group. (F) Bioluminescence imaging of NOG-hIL6 on day 8 and day 44 after a single IV administration of 10 x 10^6^ viable non-engineered or luciferase-engineered BCMs on day 1.

FIX-Padua production durability *in vivo* was assessed following a single dose of BE-101 (20×10^6^ viable cells/mouse) and monitoring plasma FIX-Padua protein levels for up to 184 days (**Figure 6A**). Following an initial decrease in levels immediately post-infusion, plasma FIX-Padua levels stabilized within 2wk and were consistently detected throughout the study (**Figure 6B**). No measurable level of FIX-Padua protein was detected in the mouse plasma collected from the vehicle control treatment group. Similarly, sustained plasma levels of human IgG (**Figure 6B**) and IgM (**Figure S2**) were detected. Human-specific FIX-Padua activity was measured in 3 mouse plasma samples using an immunocapture chromogenic activity assay from Study #2 (**Figure 6B**) with specific activity ranging from 136 IU/mg to 274 IU/mg at selected time points from day 28 to day 126.

**Figure 6.**
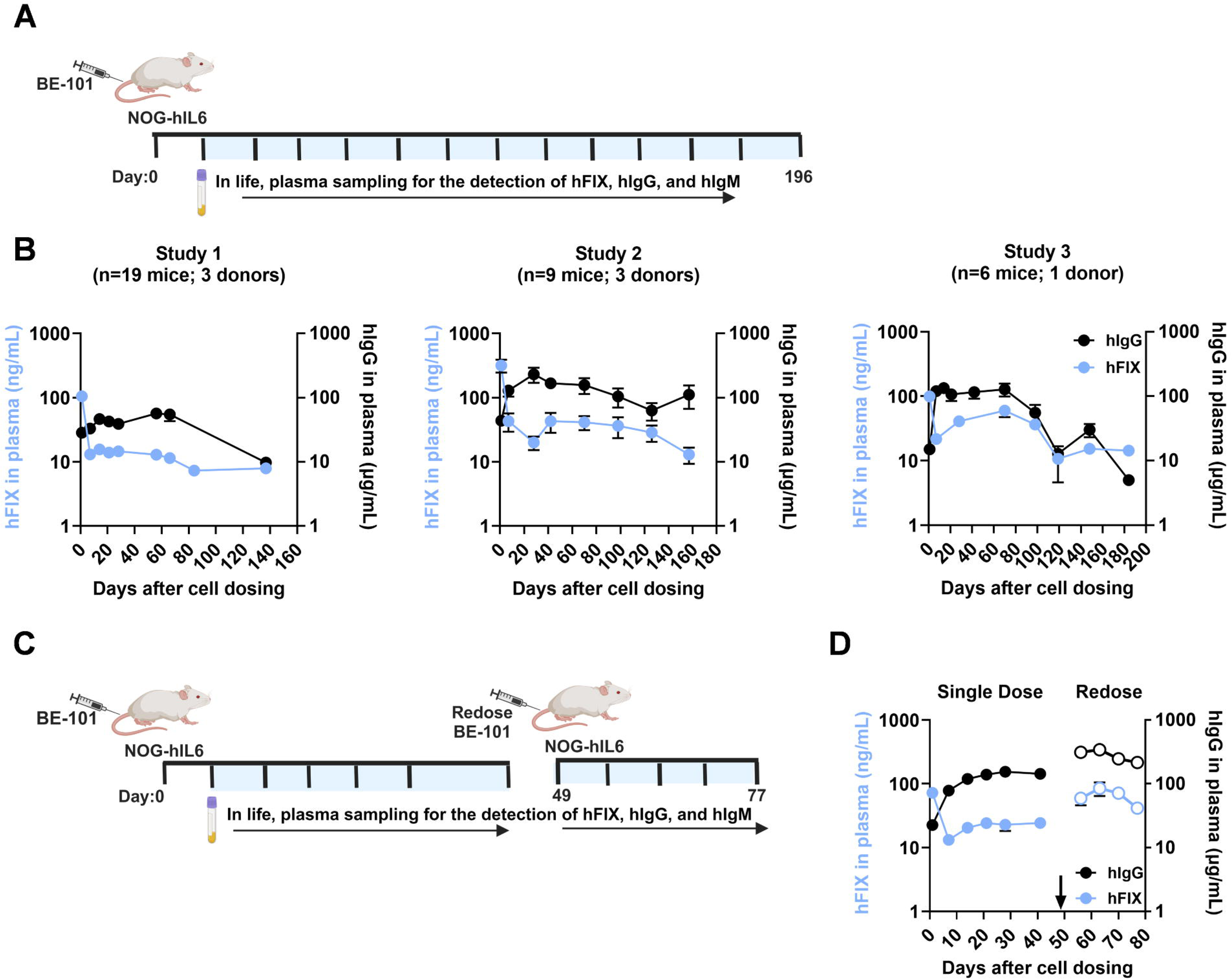
*In vivo* secretion of engineered B cell derived FIX-Padua and human IgG after a single administration and repeated administrations of BE-101 in NOG-IL6 mice. A) NOG-hIL-6 mice were dosed with 20×10^6^ total viable cells of BE-101 and endogenous B cell derived human IgG and engineered B cell derived FIX-Padua levels were measured in blood plasma up to 196 days of study. B) Data is representative of three individual studies. The lower limit of quantitation (LLOQ) of FIX-Padua protein and human IgG assay is 3.6 ng/ml and 3.4 ng/mL, respectively. C) NOG-hIL6 mice (n =4) were dosed with 20×10^6^ total viable cells of BE-101 on day 0 and re-dosed on day 49 of the study. Human IgG and FIX-Padua protein levels were measured in blood plasma after 1 day post BE-101 administration and then weekly up to 77 days of study. D) The lower limit of quantitation (LLOQ) of human FIX-Padua protein and human IgG is 2.5 ng/mL and 0.781 ng/mL, respectively. B) and D) Each data point represents the mean across all groups at that timepoint. The error bars indicate the Standard Error of the Mean (SEM).

Though direct assessment of the AAV vector copies remaining in cells at the end of the cell culture or *in vivo* was not conducted, FIX-Padua production from potential episomal AAV genomes was assessed. In research-scale studies, cells transduced in the absence of RNP (BE-101 no RNP) produced less than 4% of FIX-Padua protein compared to BE-101 control cells (**Table S6**). This was confirmed in a clinical scale process development run where cells transduced in the absence of RNP generated less than 1% of the FIX-Padua protein as compared to the average FIX-Padua protein production for clinical manufacturing runs of BE-101 where the RNP was included. These data demonstrated that the contribution of non-integrated *F9* transgene to total FIX-Padua protein production *ex vivo* was very low, and suggests that neither non-integrated *F9* transgene would contribute significantly to FIX-Padua expression, nor would they affect the potency of BE-101 *in vivo* after clinical administration.

BE-101 redosing potential in the NOG-hIL6 model was tested after a second 20×10^6^ dose of identical cryopreserved BE-101 cells from the same donor was administered on day 49 (**Figure 6C**). The mean level of FIX-Padua protein on day 56 increased 2.4-fold compared to day 42 (59 ng/mL vs. 24 ng/mL) and levels were sustained for at least 4wk (**Figure 6D**).

Similarly, mean levels of human IgG (**Figure 6D**), and IgM (**Figure S3**) on day 56 increased 2.2-3.0-fold compared to day 42 (312 μg/mL vs 143 μg/mL for human IgG, and 3.74 μg/mL vs. 1.26 μg/mL for human IgM) and were sustained for at least 4wk.

### BE-101 Safety

In a 28-day GLP toxicology study, 18 mice were treated with BE-101 manufactured from 2 donors (2.0×10^7^ total viable cells/mouse). In three long-term *in vivo* pharmacology and tolerability studies, 36 mice were treated with BE-101 manufactured from 8 donors (2.0×10^7^ total viable cells/mouse). No unscheduled deaths occurred, no abnormal clinical observations were observed, and at termination there were no adverse effects noted in clinical pathology samples or histopathology samples (**Tables S7-S13, Figure S4 and S5**).

## Discussion

Autologous, cell-based therapeutics are a promising treatment approach for many diseases.^27–34^ *Ex vivo* gene editing strategies allow precise control when modifying cellular DNA and mitigate concerns of immune responses or insertional mutagenesis in other cell types arising from *in vivo* viral vector-delivered gene therapy.^35–37^ BE-101 is a precision *ex vivo* engineered B cell medicine that produces sustained levels of highly active FIX-Padua. Terminally differentiated human PCs derived from genetically engineered B cells offer natural longevity, capacity for high levels of protein secretion, the ability to engraft without host preconditioning, and the possibility to re-dose, making them an attractive platform for the sustained supply of biologics *in vivo* where continuous dosing is required to achieve therapeutic benefit.

BE-101 exhibited robust FIX-Padua protein production *in vitro* (∼540 molecules/cell/second), greater than previous reports.^16,38^ BE-101 rapidly engrafted into mice with sustained circulation of FIX-Padua protein for up to 184 days. To our knowledge, this is the first report of *ex vivo* CRISPR/Cas9 edited, FIX-Padua-expressing PCs demonstrating stable long-term engraftment and secretion of active FIX-Padua in an animal model.

The expected BE-101 biodistribution to bone marrow tissue was confirmed in the NOG-hIL6 mouse model and BCMs were stable in marrow-containing tissues, but not other organs and tissues. Furthermore, IV administration of luciferase-expressing BCMs into NOG-hIL6 confirmed rapid homing and stability in bone marrow-harboring skeletal regions.

Although the NOG-hIL6 model is sufficient to demonstrate the durability and safety of BE-101, it does have limitations. Engraftment is estimated by the human *Alu* element biodistribution data to be less than 1% of input cells. Thus, quantifying the percentage of B cells and antibody secreting cells that contained the integrated transgene was not feasible. However, previous work on human plasma cells engineered to secrete bispecifics showed effective *in vivo* leukemia killing, suggesting edited cells stably engrafted at reproducible percentages.^39^ In humans, additional plasma cell cytokines (e.g., BAFF, APRIL), and chemokines supplied by bone marrow-resident cells such as dendritic cells, megakaryocytes, and eosinophils, as well as stromal cell extracellular matrix may support higher levels of BCM engraftment. Support for this hypothesis comes from recent studies in nonhuman primates (NHP), where we determined engraftment rates in rhesus macaque ASCs.^17^ Upon direct IV administration without preconditioning, ∼10% of the initial dose engrafted into bone marrow and ∼10% engrafted into spleen. Based on these NHP data, we projected similar levels of BE-101 engraftment would occur in human, which would be at least 20-fold higher than what was observed in the NOG-hIL6 mouse model.

Construct design and therapeutic transgene delivery are critical attributes in human gene therapy, and all aspects of BE-101 design were taken into consideration for clinical application. The FIX-Padua variant used in BE-101 was reported to be 4-8 fold more active than native FIX, a potential advantage in BCM dosing.^40–42^ Additionally, the Malmö variant, T148A, used in BE-101 construct, was the same variant used in the approved FIX therapies, Benefix®^43^ and Hemogenix®.^44^

The *CCR5* locus is minimally expressed in normal and mature B cells^45–47^ and disruption of this genomic region is not expected to affect the normal biological phenotype of B lymphocyte lineage cells. Furthermore, the *CCR5* gene has been described as a Safe Harbor Locus for stable transgene integration,^48^ and clinical evaluation of *CCR5*-edited CD4+ T cells have supported the safety of this gene editing approach in humans.^49,50^ The MND promoter was selected to drive *F9* transgene expression in BE-101 due to successful use in previous *ex vivo* gene therapy clinical trials^51,52^ and its resistance to transcriptional silencing. While prior clinical studies utilizing the MND promoter have generally exhibited a favorable safety profile, the relative strength of this promoter raises the theoretical concern of insertional mutagenesis if the site(s) of genome integration are not properly controlled as is the case for cell and gene therapies that use lentiviral and gammaretroviral vectors which promote almost random transgene insertion into the host cell genome.^53^ This potential risk of insertional mutagenesis is likely influenced by multiple factors including transduced cell type. A recent publication^54^ described the risks of insertional mutagenesis following lentiviral gene therapy in CD34+ HSCs for cerebral adrenoleukodystrophy, which appears to be associated with the inclusion of the MND promoter. However, a systemic review^55^ of T-cell lymphomas in recipients of CAR-T cells generated using lentiviral and gammaretroviral vectors concluded there is a low susceptibility of mature T cells to insertional mutagenesis and documented the almost complete lack of T-cell transformation after natural HIV infection. Moreover, the FDA estimated that a minimum of 27,000 doses of CAR-Ts, which used randomly integrating vectors containing the MND/MSCV promoters, have been infused in the United States and concluded that the risk for insertional mutagenesis was likely very low. While lentiviral gammaretroviral transduction achieves stable genomic integration in a non-targeted fashion, BE-101 utilizes targeted CRISPR/Cas9-mediated transgene insertion at the *CCR5* safe harbor locus via homology-directed repair. Moreover, we did not detect transgene insertion at the single bona-fide off-target site identified by our genotoxicity assessment. While AAV genomes are generally considered to remain episomal, random AAV integration into the genome of transduced cells have previously been described.

Unlike integrating viral vectors, random AAV vector integration frequencies are rare, with wild-type AAV integration into the known *AAVS1* insertion site estimated to be about 0.1%.^56^ In addition, recombinant AAV (rAAV) integration into the host genome is predicted to be greatly reduced because the sequences coding for the native AAV proteins needed for integration are not present in the rAAV.^57^ Importantly, no abnormal effects attributed to the administration of BE-101 were observed in long-term *in vivo* studies up to 7 months in duration. These studies included 36 total immunodeficient mice treated with BE-101 manufactured from 8 different human donors. Based on these results, we view the potential for insertional mutagenesis related to random integrations to be low.

The approach of manufacturing BE-101 using targeted CRISPR/Cas9-mediated transgene insertion at the *CCR5* locus is expected to decrease the potential for insertional mutagenesis related to random integrations observed with lentiviral or transposon-mediated transgene insertion.^55,58^ A comprehensive genotoxicity assessment was performed and submitted as part of the BE-101 regulatory filings with no issues of significance identified. Our rigorous characterization studies demonstrated that off-target effects of gCCR5_232 in BE-101 cells were limited to DNA editing rates of ∼1% at a single intergenic region over 40 kb from the nearest protein coding sequence, thus posing minimal genotoxic risk. OGM confirmed the presence of insertions conforming to HDR-mediated transgene integration at the *CCR5* on-target locus in agreement with independent ddPCR measurements. Larger insertions at this site were also detected. Although sequence level information is missing from OGM analysis, the step-wise increase in insertion size indicates potential concatemerization of viral DNA, a phenomenon observed previously in T-cell therapy^59^ and in nonclinical research.^60,61^ No other SVs above the OGM detection level were identified, indicating that genome integrity was largely preserved. In murine safety/tolerability studies (both short and long-term) there were no safety signals identified in laboratory, histopathology or mortality assessments following administration of transduced BCM cells.

The BE-101 pharmacokinetic profile is unique and distinct from what has been reported with *in vivo* FIX AAV gene therapy. Appreciable levels of FIX-Padua protein were detected within 1 day after BE-101 IV administration in mice, and steady state was reached within 2 wk. In contrast, the reported gene therapy FIX concentration-time profiles show initial FIX activity observed at ∼3 wk^9^ and steady state levels observed at 14 wk after vector infusion,^9^ requiring PwHB to maintain FIX prophylactics until steady state levels are reached. Future studies will assess whether rapid onset of FIX-Padua production with BE-101 pharmacokinetics translate to the clinic with resulting benefits to PwHB.

Equally important to the rapid onset of FIX-Padua production with BE-101 is the ability to redose/re-engraft without host pre-conditioning in NOG-hIL6 mouse model, achieving increases in FIX-Padua production in a manner close to linear dose proportionality and similar human IgG and human IgM kinetics between 1^st^ dose and repeated dose. To our knowledge, this is the first demonstration in an animal model that engineered B cells producing a human biological are redosable and titratable. These BE-101 attributes may have potential significant clinical advantages over AAV-based *in vivo* gene editing approaches in hemophilia B, where variable efficacy and toxicity may result in inadequate therapeutic windows^12^, and thus may open the possibility to re-dose.

In conclusion, BE-101, an *ex vivo* precision gene engineered BCM, produces active and sustained levels of FIX-Padua with the ability to engraft without host preconditioning. This unique biologic delivery system has the potential for redosing and titratability, and may provide a novel treatment for PwHB irrespective of age and disease severity.

## Materials and Methods

### BE-101 production

BE-101 was produced from peripheral blood B cells collected via leukapheresis from healthy human donors (StemCell Technologies, Cambridge, MA; iSpecimen, Lexington, MA; OrganaBio, South Miami, FL; and HemaCare, Lowell, MA) using commercially available selection kits (StemCell Technologies, Cambridge, MA, or Miltenyi, Charlestown, MA). Purified CD19+ B cells were cryopreserved prior to BE-101 production.

The three-phase process used for activation, expansion, and differentiation of human primary B cells and production of engineered B cells secreting therapeutic proteins have been previously described,^16,62^ and BE-101 was produced with few modifications. Isolated human B cells were thawed and cultured in Iscove’s Modified Dulbecco’s Medium (IMDM) with 10% FBS and human cytokines (Phase I media: IL-21, CpGs, and CD40L; Phase II media: IL-2, IL-6, IL-10, and IL-15; Phase III media: IFN-α2β, IL-6, and IL-15) to promote B cell differentiation into antibody secreting cells (ASCs). After activation in Phase I media for two days, human B cells were washed into electroporation buffer (Maxcyte, Gaithersburg, MD). Pre-complexed CRISPR-Cas9 and single-guide RNA (sgRNA) ribonucleoprotein (RNP) complexes (at a 2:1 guide to Cas9 ratio) were added (final concentration 4.58 μM) and the cell-RNP mixture was electroporated (B Cell Program #3). The sgRNA targeted the C-C chemokine receptor type 5 (*CCR5*) locus located at chromosome position 3p21.31. (**Table S14** has sgRNA sequence).

AAV6 vectors were packaged with a transgene encoding a human *F9* gene (hyperactive FIX-R338L, Padua variant^40–42^ and Malmö T148A variant^63^) downstream of the MND (Myeloproliferative sarcoma virus enhancer, negative control region deleted, dl587rev primer-binding site substituted) promoter,^64^ followed by a bovine growth hormone gene polyadenylation signal, and flanked by 600nt sequences homologous to genomic DNA on either side of the *CCR5* sgRNA cut site located at exon 2 of *CCR5* to mediate transgene insertion via homology-directed repair (HDR) (**Table S15** has annotated deoxyribonucleic acid (DNA) sequence of the transgene expression cassette). Following electroporation with RNP complexes, AAV6 vector was added to cells at variable multiplicities of infection and incubated for 12 to 15 minutes at 37°C followed by dilution to 1×10^6^ cells/mL in serum-free media (ExCellerate or X-VIVO 10) supplemented with Phase I cytokines, then expanded and differentiated towards the PC lineage during the remainder of the three-phase process. Control cells were cultured similarly but without the addition of AAV6 vector.

The final BE-101 drug product was a sterile cell suspension. The cryopreserved preparation, in equal quantities of Plasma-Lyte A with 5% Human Serum Albumin (HSA) and CryoStor® CS10, is identical to the product to be used in clinical trials.

### BE-101 product characterization

Characterization of BE-101 included assessment of viability, cellular immunophenotype, on-target insertion and deletion (INDEL), and transgene integration. Cell counts and viability were determined using Cellaca^MX^ (Nexcelom Bioscience, Lawrence, MA) or NC200^TM^ (ChemoMetec A/S, Denmark).

Cellular immunophenotyping was measured by flow cytometry. Cells suspended in 100 μL PBS in 96-well plates or 1.5mL Eppendorf tubes received viability dye (100 μL) and were incubated at 4°C for 10 to 15 minutes followed by washing 2x with flow cytometry staining buffer (Invitrogen, Waltham, MA) and pelleting at 450xg for 3 minutes. Following live/dead staining, surface staining antibodies were added according to manufacturers’ recommendation with specific fluorochrome-conjugated monoclonal antibodies APC-CD19, FITC-CD20, PECY7-CD27, BV421-CD38, PE-CD138 (Becton Dickinson, Franklin Lakes, NJ, and Biolegend, San Diego, CA) and incubated for 25 to 30min at 4°C. Cells were washed and pelleted and pellets were resuspended in 150μL of flow cytometry staining buffer to assess live singlet cells and B cell differentiation using the NovoCyte Penteon Flow Cytometer (Agilent Technologies, Santa Clara, CA). Over 10,000 events per sample were collected and analyzed using FlowJo software (BD Bioscience, Franklin Lakes, NJ).

INDEL analysis at the on-target sgRNA cut site was performed as described previously.^65^ (**Table S16** has sequences of primers designed via the IDT PrimerQuest tool). Transgene integration frequency was measured by Droplet Digital PCR (ddPCR) (Bio-Rad Laboratories, Hercules, CA) using isolated gDNA from the final BE-101 product. ddPCR probes were designed in house (**Table S17** has sequences of designed ddPCR probes). Transgene integration quantification was calculated as: copies/μL FAM+ droplets (target amplicon or engineered alleles)/copies/μL HEX+ droplets (reference amplicon or total alleles) * 100=%Targeted integration. (**Table S17** has primers and probes).

### Measurement of FIX-Padua secreted by engineered B cells *in vitro* and *in vivo*

FIX-Padua secretion *in vitro* was quantified by ELISA (Human Factor IX ELISA Kit, Abcam, Waltham, MA). *In vivo* analysis of plasma FIX-Padua was performed by ELISA following a selective human FIX protein capture step using human FIX monoclonal antibodies GMA102 and GMA184 (Green Mountain Antibodies, Burlington, VT) prior to determination of human FIX-Padua only in mouse plasma. FIX-Padua and Malmö variant did not interfere with the human FIX selective capture antibodies used to detect and measure the circulating human FIX-Padua protein derived from BE-101 in NOG-hIL6 mouse plasma. Activity of FIX-Padua was assessed using a chromogenic assay (Rox Factor IX chromogenic activity kit, Diapharma, West Chester, OH) following the selective human FIX protein capture step. FIX-Padua activity was also assessed in an aPTT assay following purification from cell culture supernatants using size-exclusion chromatography-ultraviolet (SEC-UV) (performed at NovaBioAssays Woburn, MA). Specific activity was calculated based on measured protein activity divided by calculated amount of protein present in the sample. Corresponding clotting times were converted to percent activity using a log-log regression; percent activity, IU/mL and IU/mg are reported.

### Assessment of DNA editing at potential gCCR5_232 off-target sites in BE-101

On- and off-target sites identified by CRISPRitz, G-GUIDE, and SITE-seq were amplified from genomic DNA of five batches of BE-101 cells and donor-matched unengineered control cells using multiplexed amplicon sequencing (rhAmpSeq, IDT, Coralville, IA) or standard amplicon sequencing. Sequencing libraries were generated according to the manufacturer’s instructions and sequenced in an Illumina MiSeq targeting a sequencing depth of ≥5000 reads per amplicon. INDEL analyses were conducted using CRISPAltRations.^66^ For all sites reaching the targeted sequencing depth, an effect-size and p-value were calculated by fitting a negative binomial generalized linear model. Statistically validated off-target sites were defined as those with estimated differential editing above 0.2% and a nominal p-value below 0.05 in a one-sided Z-test.^67^

### Evaluation of genome integrity by Optical Genome Mapping (OGM)

Sample preparation and data acquisition for OGM (Bionano Genomics) were performed by Boston Children’s Hospital IDDRC Molecular Genetics Core Facility (Boston, MA). Ultra-high molecular weight (UHMW) gDNA from three lots of BE-101 cells and donor-matched unengineered controls was extracted and labeled, loaded on the flow cell Saphyr G3.3 chip and imaged on the Saphyr instrument. Rare Variant Analysis (RVA), *De Novo* Assembly, and variant calling were executed on Bionano Solve software (v3.8.1). Reporting and visualization of structural variants (SVs) was done on Bionano Access (v1.8.1) using hg38 as reference genome. Each engineered sample was compared against its donor-matched unengineered control. With 300X coverage, Bionano can detect SVs with 5% events and with over 90% sensitivity.

### Animal studies for BE-101 engraftment, biodistribution, and persistence

Immunodeficient NOG-hIL6 mice (*NOD.Cg-Prkdcscid Il2rgtm1SugTg(CMV-IL6)1-1Jic/JicTac*, Taconic Biosciences Rensselaer, NY) engineered to express a human interleukin 6 (hIL-6) transgene,^68^ 8–14-week-old, were housed (up to 3 animals per cage) in a HEPA filtered, individually isolated cage system (Innovive, San Diego, CA) and acclimated for at least 3d. Housing was an American Association for Accreditation of Laboratory Animal Care (AAALAC) accredited vivarium operated by SmartLabs (Cambridge, MA); all animal care and use procedures were approved by the SmartLabs Institute Animal Care and Use Committee (IACUC) prior to study start. Mice were randomly assigned to study groups.

A BE-101 surrogate construct expressing firefly luciferase was used to assess biodistribution and persistence. Surrogate cells formulated in PBS buffer at a concentration of 5×10^7^ cells/mL were injected intravenously at 1×10^7^ viable cells/mouse. Cryo-preserved and thawed BE-101 cells were formulated in PlasmaLyte or PBS buffer at a concentration of 1×10^8^ cells/mL and injected intravenously at 2×10^7^ viable cells/mouse. For redosing, cryopreserved BE-101 cells were dosed at 2×10^7^ total viable cells/mouse on study days 0 and 49. Animals were observed twice per week for changes in body condition and weights. Blood samples were collected by tail bleeding into Microvette CB300 EDTA K2E containers (Sarstedt, Nümbrecht, Germany). Plasma FIX-Padua protein level and immunocapture chromogenic activity were measured as described above. At study end, mice were euthanized, and blood and tissue samples were collected for biodistribution, clinical chemistry, or histopathology assessments.

Supplemental information includes methodology on LC-MS/MS analysis of post-translational modification of the GLA domain in BE-101-derived FIX-Padua protein, nomination of potential off-target sites (i.e., CRISPRitz, G-GUIDE, and SITE-seq), human selective IgG and IgM measurements in mouse plasma, *Alu* PCR assay assessing BE-101 biodistribution, whole body bioluminescence of a BE-101 surrogate construct expressing firefly luciferase dosed to NOG-hIL6 mice, and safety based on 28 day GLP toxicology and long-term *in vivo* pharmacology and tolerability studies in NOG-hIL6 mice.

## Data Sharing Statement

All relevant data are included in the manuscript. Any additional data is available on request from the corresponding author, Hanlan Liu at.

## Acknowledgments

The authors thank Dr. David Rawlings and Dr. Richard James at Seattle Children’s Research Institute, Dr. Glenn Pierce at the World Federation of Hemophilia, Dr. Wing Yen Wong and Dr. Krishnan Viswanadhan at Be Biopharma for reviewing and providing valuable inputs to the manuscript.

The authors thank Technical Development Team at Be Biopharma for generating BE-101 materials for in vivo studies and genotoxicity assessments. The authors also thank Daniel Ferguson for helping some of the BE-101 genotoxicity characterization. Patrice C. Ferriola, PhD, of KZE PharmAssociates, LLC aided in preparation of the manuscript and was funded by Be Biopharma.

## Authorship Contributions

H.L. S.S., T.J.M., C.B., S.K., T.P., S.T., A. Lundberg, R.C., S.T., C.Y., S.D., L.L. W.B., E.L., C.R.M., S.C., R.K., A.F.H., S.A., A. Lazorchak, and C.S. designed and/or performed experiments; S.S., T.O., E.L. and H-M.C. performed computational data analysis; H.L., S.S., T.M., S.K., C.L., H-M.C., A.F.H., S.A., A. Lazorchak, and R.A.M. wrote the manuscript.

## Declaration of interests

H.L. S.S., T.J.M., C.B., S.K., S.T., A. Lundberg, R.C., S.D., L.L. W.B., E.L., C.R.L., H-M.C., R.K., A.F.H., S.A., A. Lazorchak, and R.A.M. reported being current employees at Be Biopharma. H.L. S.S., T.J.M., C.B., S.K., T.P., S.T., A. Lundberg, R.C., S.T., C.Y., S.D., L.L. W.B., E.L., C.R.M, C.R.L., T.K.O., S.C., H-M.C., R.K., A.F.H., S.A., A. Lazorchak, and R.A.M. reported holding stock options of Be Biopharma, a privately-held company. The remaining authors declare no competing financial interests. T.J.M, H.L., A.F.H, S.A., S.S., S.K., and R.A.M. are co-inventors of the patent “Engineered cell preparations for treatment of hemophilia” (WO2024220446A2).

