## Supplemental Materials for "A Precision Gene Engineered B Cell Medicine Producing Sustained Levels of Active Factor IX for Hemophilia B Therapy"

**Supplemental Data**

**Supplemental Methods**

**Mass spectrometric characterization of FIX derived from BE-101**

Mass spectrometric sample preparation, methanolic HCl derivatization to detect γ-carboxyglutamic acid in peptides by positive ion mode LC-ESI-MS/MS, and mass spectrometric characterization of purified FIX protein derived from BE-101 cultured in the presence of vitamin K3 followed the procedures as previously described[1] with minor modification.

**Nomination of potential off-target sites**

**In silico prediction.** Computational prediction of potential off-target sites in GRCh38 with homology to the intended target site of gCCR5_232 (with no more than four nucleotide mismatches and two bulges, CFD score ≥0.2,[2] and NGG PAM sequence) was performed using CRISPRitz v2.5.8.[3]

**G-GUIDE.** Primary human B cells from five unique donors were transfected with Cas9–sgRNA RNP complexes and end-protected dsODNs via electroporation on the MaxCyte ATx™ using B Cell Program #3. Transfected cells were cultured for 48h and collected for gDNA extraction. The G-GUIDE™ assay was provided and conducted by GeneGoCell Inc (San Diego, CA). DNA was sheared by sonication to an average length of 500 bp, end-repaired, A-tailed and ligated to Y-shape adapter. The adapter-ligated library was subjected to three rounds of nested PCR using dsODN sense- and antisense-specific primers in separate reactions. Libraries were quantified on a Qubit Fluorometer 4 (Thermo Fisher Scientific, Waltham, MA). Library size was measured on a TapeStation 4200 (Agilent Technologies, Santa Clara, CA). Successful libraries were then sequenced on NextSeq 2000 (Illumina, San Diego, CA). GUIDE analysis was performed as previously described.[4] Nomination of genomic loci as potential off-targets required the left and right side of the dsODN tag integration site to be detected in both technical replicates in at least one B cell donor.

**SITE-seq.** SITE-seq was performed as described previously with minor modifications.[29] Wild-type high molecular weight (HMW) genomic DNA (gDNA) was purified from B cells using Monarch HMW DNA extraction kit for cells and blood (NEB). In vitro cleavage reactions were performed using 10 µg of gDNA from six unique B cell donors, with 4.58 µM of Cas9 and two-fold molar excess of GMP-grade gCCR5_232 in Cas9 buffer (20mM HEPES, 0.1M NaCl, 5mM $MgCl_2$, 0.1mM EDTA). Cleaved products were A-tailed, ligated with an Illumina-compatible biotinylated adapter, and treated with dsDNA fragmentase (NEB) to generate DNA fragments of 800-bp on average. Fragmented DNA was end-repaired, and a second Illumina-compatible adapter was ligated. Dual adapter-ligated DNA molecules were purified using M-280 Dynabead (Thermo Fisher, Waltham, MA) and amplified by PCR with barcoded universal primers. Libraries were quantified by qPCR and sequenced with 150-bp paired-end reads on an Illumina MiSeq. Calling Cas9 cleavage sites required ≥5 reads terminating at the same nucleotide, and a sequence with no more than 7 mismatches and 5 bulges relative to the intended target site of gCCR5_232 to be found in a +/-35 bp window around the cut site. Sites were nominated as potential off-targets if found in all three technical replicates of at least two out of six profiled donors.

**Human selective IgG and IgM measurement in mouse plasma**

Human selective IgG and IgM measurement in mouse plasma was determined by either ELISA or MSD immunoassay. For ELISA assay, briefly, 384 well maxisorp plates were coated with capture antibody, anti-human IgG (Southern Biotech, Birmingham, AL, Cat# 9040-01) at 3 μg/mL or anti-human IgM (Southern Biotech, Birmingham, AL, Cat# 9022-01) at 1 μg/mL in carb-bicarb buffer, overnight at 4 °C then plates were washed 3 times with an automated plate washer. The antibody coated plates were blocked against non-specific binding with 1% BSA for 1 hour at room temperature. Plasma samples or standards were added to the wells of the plate and incubated for 1.5 hours at room temperature. Plates were washed as previously described. Plate-bound hIgG or plate-bound hIgM was then captured by secondary antibody, goat anti-IgG polyclonal antibody HPR (Southern Biotech, Birmingham, AL, Cat# 2014-05) or goat anti-IgM polyclonal antibody HPR (Southern Biotech, Birmingham, AL, Cat# 2023-05) at 1:5000 dilution for 1 hour at room temperature. TMB two-part substrate was added for 1 min (development time to be determined empirically) and then the reaction was stopped. The plate absorbance was read at 450 nm and 570 nm by a Cytation5 Plate Reader. Sample values were interpolated using the standards at known human IgG or human IgM concentrations. MSD immunoassay plates were precoated with capture IgG and IgM antibodies on independent and well-defined spots. Next, the MSD plates were first blocked using 1X PBS with 1% BSA blocking buffer, prior to the addition of calibration standards, quality control samples, and test samples. Mouse plasma or serum samples were diluted in diluent 100 to a minimum of 64-fold before adding to the MSD plate. All samples were incubated for 2 hours at room temperature, shaking at 700 rotations per minute (rpm), then the plate was washed, and sulfo-tag labelled anti-human/non-human primate (NHP) detection antibodies (Meso Scale Diagnostics, LLC,

Rockville, MD, Cat#D20JL-6) were added to all wells at 1:50 dilution in diluent 100 and incubated for two hours at room temperature, shaking at 700 rpm. Analytes in sample bound to capture antibodies immobilized on the working electrode surface; recruitment of the detection antibodies by the bound analytes completed the sandwich. After another wash, read buffer was added to all wells, which provided an appropriate chemical environment for electrochemiluminescence. The plate was loaded onto the instrument, where voltage was applied to the plate electrodes and caused captured labels to emit light. The instrument measured the intensity of emitted light to provide quantitative measure of analytes in the sample. Human IgG and IgM concentrations were calculated by five parameter logistic regression model to appropriate reference protein standards.

***Alu* PCR assay assessing BE-101 biodistribution**

Quantitation of human genomic DNA (hgDNA) by detecting *Alu* in NOG-IL6 mouse gDNA was analyzed by a qPCR assay using an ABI 7 Pro system (ThermoFisher Scientific, Waltham, MA). From the qualification of the method, an eight-point standard curve was established using serial dilutions of extracted DNA of control B cell line ranging from $1.00 \times 10^6$ to 50.0 fg of hgDNA/rxn in water and evaluated in the presence of 100 ng of mouse genomic DNA (mgDNA) in the qPCR reaction. These reference standards were prepared for the calibration curve and were examined in six independent qPCR runs to set up suggested assay criteria before sample analysis . The LOD for this method was set at 50.0 fg of hgDNA/100 ng of mgDNA and derived from the qualification runs. In addition, the LLOQ of the qPCR assay was also determined from the qualification runs and was set to be 100 fg of hgDNA/rxn by spiking defined reference standard curve hgDNA in the 100 ng of mouse kidney extracted gDNA matrix. The qPCR assay included positive and negative controls and could quantify ≥ 100 fg of hgDNA/100 ng of mgDNA. Similarly, the ULOQ ($1.00 \times 10^6$ fg of hgDNA/ 100 ng of mgDNA) was also determined in method qualification runs.

**Whole body bioluminescence of a BE-101 surrogate construct expressing firefly luciferase dosed into NOG-hIL6 mice**

For bioluminescence imaging, the PerkinElmer in vivo imaging system (IVIS) was used to monitor the engraftment and bio-persistence of adoptively transferred human B cells that have been engineered to express firefly luciferase. Prior to bioluminescent imaging, the mice were dosed with 200 μL (150 mg/kg) of RediJect D-Luciferin by subcutaneous (SC) injection. The mice were anesthetized in an isoflurane induction chamber prior to being moved into the IVIS system where they were maintained under isoflurane anesthesia via a nose cone manifold. This manifold allows up to five mice to be imaged at one time. A heated stage maintains the body temperature of the animals during image capture and mice can remain within the IVIS for up to 40 min. Upon being removed from the IVIS, the mice were observed until they fully regained consciousness. The optimal time window for imaging was determined empirically by luciferase kinetic analysis in the mouse test system.

Intraperitoneal dosing of D-Luciferin: The aim of this technique was to administer material into the space between the abdominal organs and the internal wall of the abdomen, avoiding injection directly into an organ. To prevent accidental puncture of organs, mice were restrained and held with their ventrum exposed and head pointed downward. The injection site was wiped with a sterile alcohol pad. A 25 gauge or smaller needle was inserted at a 40-degree angle into the abdominal cavity to the left of the abdominal midline in the lower left quadrant of the animal, to avoid the cecum and urinary bladder. Prior to injection, the syringe was aspirated to ensure that a vein had not been entered accidentally.

**Safety of FIX-Padua-expressing BE-101 in 28-day GLP toxicology and long-term *in vivo* pharmacology and tolerability studies in NOG-hIL6 mice**

A 28-day single-dose GLP-compliant toxicology study was conducted in NOG-hIL6 mice. Plasma levels of hFIX in this model peaked the day after administration and generally stabilized by 4 weeks after a single administration of BE-101. Since the B lymphocyte lineage cells do not expand after administration, and no changes in pharmacology occur after the 4-week timepoint, a study duration of 4 weeks was considered sufficient for assessing any potential acute toxicity from BE-101. Material for the study GLP study was generated from two independent healthy human donors and 9 male mice per donor were administered a single dose of BE-101. A vehicle group of 15 male mice were administered plasmalyte only. Engraftment of BE-101 was confirmed by measuring hFIX in the plasma 21 days after administration and confirming the presence of human DNA in the bone marrow at the end of the study. There were no abnormal clinical observations associated with BE-101 treatment, no significant differences in body weight between treatment groups, and all animals survived until their scheduled termination. At termination, clinical pathology assessments showed no significant differences between groups and no BE-101-related effects on coagulation parameters (fibrinogen, PT, and APTT), hematology parameters (complete blood counts), or clinical chemistry parameters (22 serum chemistry assays). There were no BE-101 associated findings at necropsy and no microscopic findings considered to be related to BE-101 in a comprehensive set of tissues assessed from all animals that received BE-101.

Since BE-101 persists in the NOG-hIL6 model beyond 28 days, long term safety was also assessed in 3 longer term pharmacology studies in NOG-hIL6 mice. In a 21-week pharmacology study, a total of 21 female mice were administered BE-101 with no BE-101-related mortality, abnormal clinical observations, or abnormalities identified at necropsy. In a 22-week pharmacology study, a total of 9 female mice were administered BE-101 and there were no unscheduled deaths or abnormal observations and no significant differences in body weights. At termination, there were no gross abnormalities noted and histopathological examination of 8 tissues per animals determined that there were no microscopic findings associated with BE-101 treatment, nor were differences seen between animals receiving cells from different donors. In a 28-week pharmacology study, 6 female mice were administered BE-101 and there was no mortality, abnormal clinical observations, or abnormalities at necropsy. Limited hematology and clinical chemistry samples at necropsy were normal, and there were no microscopic findings in the 20 tissues analyzed per animal. Overall, the totality of safety data from long term studies up to 7 months in duration includes 36 total mice treated with BE-101 ($2 \times 10^7$ total viable cells/mouse) manufactured from 8 independent donors, without any abnormal effects attributed to the administration of BE-101.

### Supplemental Table

### Table S1: Phenotypic Analysis of Leukocyte Populations in BE-101 Drug Product

| **Population** | | **Batch ID** | | | |
|---|---|---|---|---|---|
| | | **BTD23087** | **BTD23092** | **BTD23118** | **2208_102-2023_DP** |
| T Cells | CD45+CD3+CD16-CD56- | 0.3 % | 0.4 % | 0.5 % | 0.1 % |
| Monocytes | CD45+CD14+ | 0.1 % | 0.1 % | 0.1 % | 0.4 % |
| NK Cells | CD45+CD3-CD16+CD56+ | 0.3 % | 0.8 % | 0.6 % | 0.5 % |
| NKT Cells | CD45+CD3+CD16+CD56+ | 0.3 % | 0.4 % | 0.5 % | 0.1 % |
| Sum of Non-B Lineage Cells * | | 1.0 % | 1.7 % | 1.7 % | 1.2 % |
| Sum of all B lymphocyte lineage cells | CD45+CD3-CD16-CD56- CD38+ and/or CD19+ | 98.8 % | 97.9 % | 98.1 % | 98.5 % |

* sum of % values above

### Table S2: Phenotypic Analysis of B Lymphocyte Lineage Cells in BE-101 Drug Product

| Population | BTD23087 | BTD23092 | BTD23118 | 2208_102-2023_DP |
|---|---|---|---|---|
| CD38+ | 94.5% | 94.7% | 92.3% | 95.3% |
| CD27+ | 87.6% | 80.0% | 82.7% | 94.9% |
| CD38+ CD27+ | 86.1% | 79.3% | 82.0% | 93.0% |
| CD38+CD27+CD138+ | 23.5% | 19.2% | 17.2% | 31.2% |
| CD38+CD27+CD138- | 62.6% | 60.1% | 64.8% | 61.8% |
| CD19+ | 96.0% | 96.7% | 98.6% | N/A |
| CD20+ | 9.0% | 19.1% | 28.8% | N/A |
| IgM+ | 27.1% | 22.4% | 28.5% | N/A |
| IgD+ | 50.0% | 40.2% | 53.9% | N/A |

Gating based on live, singlet cells, setting live cells as 100%. CD45 not included in this panel.

**Table S3: Off-target sites identified by G-Guide, SITE-Seq, and CRISPRitz**

| Cut Position | Genome Region | G-GUIDE | SITE-Seq | CRISPRitz |
|---|---|---|---|---|
| **chr3:46373214:+** | CDS | TRUE | TRUE | TRUE |
| **chr5:32668418:+** | INTERGENIC | TRUE | TRUE | TRUE |
| **chr12:12823395:+** | INTRON | TRUE | TRUE | TRUE |
| **chr1:204082783:+** | INTRON | FALSE | TRUE | TRUE |
| **chr11:83710971:+** | INTRON | FALSE | TRUE | TRUE |
| **chr12:28362143:-** | INTRON | FALSE | TRUE | TRUE |
| **chr12:33025504:+** | INTERGENIC | FALSE | TRUE | TRUE |
| **chr12:41709634:+** | INTERGENIC | FALSE | TRUE | TRUE |
| **chr13:74757144:-** | INTERGENIC | FALSE | TRUE | TRUE |
| **chr14:72115118:+** | INTRON | FALSE | TRUE | TRUE |
| **chr17:16011638:-** | INTRON | FALSE | TRUE | TRUE |
| **chr2:37650518:+** | INTRON | FALSE | TRUE | TRUE |
| **chr3:154271332:-** | INTERGENIC | FALSE | TRUE | TRUE |
| **chr4:58670079:-** | INTERGENIC | FALSE | TRUE | TRUE |
| **chr5:156265830:+** | INTRON | FALSE | TRUE | TRUE |
| **chr5:63564110:-** | INTERGENIC | FALSE | TRUE | TRUE |
| **chr7:95775934:+** | INTRON | FALSE | TRUE | TRUE |
| **chr8:25536028:-** | INTERGENIC | FALSE | TRUE | TRUE |
| **chr8:34129360:-** | INTERGENIC | FALSE | TRUE | TRUE |
| **chr8:52307651:-** | INTRON | FALSE | TRUE | TRUE |
| **chr8:96106077:-** | INTERGENIC | FALSE | TRUE | TRUE |
| **chr3:48263061:-** | INTRON | TRUE | FALSE | TRUE |
| **chr1:110013819:+** | INTRON | FALSE | TRUE | FALSE |
| **chr1:112048261:+** | INTERGENIC | FALSE | TRUE | FALSE |
| **chr1:117536366:-** | INTERGENIC | FALSE | TRUE | FALSE |
| **chr1:117713321:-** | INTERGENIC | FALSE | TRUE | FALSE |
| **chr1:173339902:+** | INTERGENIC | FALSE | TRUE | FALSE |
| **chr1:192001446:+** | INTRON | FALSE | TRUE | FALSE |
| **chr1:215080829:+** | INTRON | FALSE | TRUE | FALSE |
| **chr1:45215920:+** | INTRON | FALSE | TRUE | FALSE |
| **chr1:50529104:+** | INTRON | FALSE | TRUE | FALSE |
| **chr1:72852997:-** | INTRON | FALSE | TRUE | FALSE |
| **chr1:80448623:+** | INTERGENIC | FALSE | TRUE | FALSE |
| **chr1:88339536:+** | INTRON | FALSE | TRUE | FALSE |
| **chr10:101149488:+** | INTERGENIC | FALSE | TRUE | FALSE |
| **chr10:60336288:+** | INTRON | FALSE | TRUE | FALSE |
| **chr10:68386104:+** | CDS | FALSE | TRUE | FALSE |
| **chr11:130323657:+** | INTRON | FALSE | TRUE | FALSE |

| | | | | |
|---|---|---|---|---|
| **chr11:131372205:+** | INTRON | FALSE | TRUE | FALSE |
| **chr11:54592486:+** | INTERGENIC | FALSE | TRUE | FALSE |
| **chr11:55117942:+** | INTERGENIC | FALSE | TRUE | FALSE |
| **chr11:79267307:+** | INTRON | FALSE | TRUE | FALSE |
| **chr11:91283839:+** | INTERGENIC | FALSE | TRUE | FALSE |
| **chr11:91662198:+** | INTERGENIC | FALSE | TRUE | FALSE |
| **chr12:130937345:-** | INTERGENIC | FALSE | TRUE | FALSE |
| **chr12:65538087:+** | INTRON | FALSE | TRUE | FALSE |
| **chr12:67099738:-** | INTERGENIC | FALSE | TRUE | FALSE |
| **chr12:76846031:+** | INTRON | FALSE | TRUE | FALSE |
| **chr13:25514926:+** | INTRON | FALSE | TRUE | FALSE |
| **chr13:66509424:-** | INTRON | FALSE | TRUE | FALSE |
| **chr13:84828188:-** | INTERGENIC | FALSE | TRUE | FALSE |
| **chr13:95893936:+** | INTRON | FALSE | TRUE | FALSE |
| **chr14:21313851:+** | INTRON | FALSE | TRUE | FALSE |
| **chr14:67173277:-** | INTRON | FALSE | TRUE | FALSE |
| **chr14:90250368:+** | INTERGENIC | FALSE | TRUE | FALSE |
| **chr15:101681301:+** | INTRON | FALSE | TRUE | FALSE |
| **chr15:43955169:+** | INTRON | FALSE | TRUE | FALSE |
| **chr15:71184426:-** | INTRON | FALSE | TRUE | FALSE |
| **chr15:86521385:+** | INTRON | FALSE | TRUE | FALSE |
| **chr15:95330154:+** | INTRON | FALSE | TRUE | FALSE |
| **chr16:7473048:+** | INTRON | FALSE | TRUE | FALSE |
| **chr16:7515001:-** | INTRON | FALSE | TRUE | FALSE |
| **chr17:58165113:-** | INTERGENIC | FALSE | TRUE | FALSE |
| **chr17:66458937:+** | INTRON | FALSE | TRUE | FALSE |
| **chr18:109193:+** | INTERGENIC | FALSE | TRUE | FALSE |
| **chr18:38872155:+** | INTERGENIC | FALSE | TRUE | FALSE |
| **chr18:62986538:-** | INTERGENIC | FALSE | TRUE | FALSE |
| **chr18:76210758:-** | INTRON | FALSE | TRUE | FALSE |
| **chr19:30620679:+** | INTRON | FALSE | TRUE | FALSE |
| **chr2:12375207:+** | INTRON | FALSE | TRUE | FALSE |
| **chr2:12415399:+** | INTRON | FALSE | TRUE | FALSE |
| **chr2:178945257:+** | INTRON | FALSE | TRUE | FALSE |
| **chr2:190484162:-** | INTRON | FALSE | TRUE | FALSE |
| **chr2:19190082:+** | INTRON | FALSE | TRUE | FALSE |
| **chr2:46465105:-** | INTRON | FALSE | TRUE | FALSE |
| **chr2:70566588:+** | INTERGENIC | FALSE | TRUE | FALSE |
| **chr2:79919202:+** | INTRON | FALSE | TRUE | FALSE |
| **chr3:1107171:-** | INTRON | FALSE | TRUE | FALSE |
| **chr3:120175975:+** | INTRON | FALSE | TRUE | FALSE |

| | | | | |
|---|---|---|---|---|
| **chr3:146458072:+** | INTRON | FALSE | TRUE | FALSE |
| **chr3:158063485:+** | INTRON | FALSE | TRUE | FALSE |
| **chr3:178348514:+** | INTRON | FALSE | TRUE | FALSE |
| **chr3:62514904:-** | INTRON | FALSE | TRUE | FALSE |
| **chr3:64724730:+** | INTRON | FALSE | TRUE | FALSE |
| **chr3:66875863:+** | INTERGENIC | FALSE | TRUE | FALSE |
| **chr3:84551563:+** | INTERGENIC | FALSE | TRUE | FALSE |
| **chr4:107551526:+** | INTERGENIC | FALSE | TRUE | FALSE |
| **chr4:120834595:+** | INTRON | FALSE | TRUE | FALSE |
| **chr4:160878787:+** | INTERGENIC | FALSE | TRUE | FALSE |
| **chr4:88218179:-** | INTRON | FALSE | TRUE | FALSE |
| **chr4:90969269:+** | INTRON | FALSE | TRUE | FALSE |
| **chr5:139706319:+** | INTRON | FALSE | TRUE | FALSE |
| **chr5:16981253:-** | INTERGENIC | FALSE | TRUE | FALSE |
| **chr5:32025982:-** | INTRON | FALSE | TRUE | FALSE |
| **chr5:57042990:+** | INTERGENIC | FALSE | TRUE | FALSE |
| **chr5:65763770:-** | INTRON | FALSE | TRUE | FALSE |
| **chr5:96353436:+** | INTRON | FALSE | TRUE | FALSE |
| **chr6:144603896:+** | INTRON | FALSE | TRUE | FALSE |
| **chr6:71308252:-** | UTR | FALSE | TRUE | FALSE |
| **chr6:8196913:-** | INTRON | FALSE | TRUE | FALSE |
| **chr7:117265746:-** | INTERGENIC | FALSE | TRUE | FALSE |
| **chr7:117268616:-** | INTERGENIC | FALSE | TRUE | FALSE |
| **chr7:120141414:+** | INTRON | FALSE | TRUE | FALSE |
| **chr7:34326474:-** | INTERGENIC | FALSE | TRUE | FALSE |
| **chr7:69712882:-** | INTRON | FALSE | TRUE | FALSE |
| **chr7:87131213:+** | INTERGENIC | FALSE | TRUE | FALSE |
| **chr8:130408203:+** | INTRON | FALSE | TRUE | FALSE |
| **chr8:14490205:+** | INTRON | FALSE | TRUE | FALSE |
| **chr8:16391992:+** | INTRON | FALSE | TRUE | FALSE |
| **chr8:18583516:+** | INTRON | FALSE | TRUE | FALSE |
| **chr8:71234842:+** | INTRON | FALSE | TRUE | FALSE |
| **chr9:15937891:-** | INTRON | FALSE | TRUE | FALSE |
| **chr9:17100115:-** | INTRON | FALSE | TRUE | FALSE |
| **chr9:6566360:+** | INTRON | FALSE | TRUE | FALSE |
| **chr9:96441610:+** | INTERGENIC | FALSE | TRUE | FALSE |
| **chr9:98446972:+** | INTRON | FALSE | TRUE | FALSE |
| **chrX:108952195:+** | INTERGENIC | FALSE | TRUE | FALSE |
| **chrX:121844185:-** | INTERGENIC | FALSE | TRUE | FALSE |
| **chrX:94296164:+** | INTERGENIC | FALSE | TRUE | FALSE |
| **chrY:11312114:-** | INTERGENIC | FALSE | TRUE | FALSE |

| | | | | |
|---|---|---|---|---|
| **chrY:14296934:+** | INTERGENIC | FALSE | TRUE | FALSE |
| **chr1:125180347:+** | INTERGENIC | TRUE | FALSE | FALSE |
| **chr14:105859522:+** | INTERGENIC | TRUE | FALSE | FALSE |
| **chr14:105859686:-** | INTERGENIC | TRUE | FALSE | FALSE |
| **chr3:46373168:+** | CDS | TRUE | FALSE | FALSE |
| **chr4:49108432:+** | INTERGENIC | TRUE | FALSE | FALSE |
| **chr1:7491434:-** | INTRON | FALSE | FALSE | TRUE |
| **chr5:106088152:+** | INTERGENIC | FALSE | FALSE | TRUE |
| **chr5:150021173:-** | INTRON | FALSE | FALSE | TRUE |
| **chr5:3465800:-** | INTRON | FALSE | FALSE | TRUE |
| **chr6:160922994:+** | INTRON | FALSE | FALSE | TRUE |
| **chr5:14756639:-** | INTRON | FALSE | FALSE | TRUE |
| **chr6:50514343:-** | INTRON | FALSE | FALSE | TRUE |
| **chr5:14347823:+** | INTRON | FALSE | FALSE | TRUE |
| **chr12:84481668:-** | INTERGENIC | FALSE | FALSE | TRUE |
| **chr16:57149878:+** | INTERGENIC | FALSE | FALSE | TRUE |
| **chr6:77475446:-** | INTERGENIC | FALSE | FALSE | TRUE |
| **chr10:17188356:-** | INTRON | FALSE | FALSE | TRUE |
| **chr20:15626418:+** | INTRON | FALSE | FALSE | TRUE |
| **chr10:4663235:+** | INTRON | FALSE | FALSE | TRUE |
| **chr7:124507009:-** | INTERGENIC | FALSE | FALSE | TRUE |
| **chr3:85474768:+** | INTRON | FALSE | FALSE | TRUE |
| **chr3:176012031:+** | INTERGENIC | FALSE | FALSE | TRUE |
| **chr3:3922827:+** | INTRON | FALSE | FALSE | TRUE |
| **chr3:84394873:-** | INTERGENIC | FALSE | FALSE | TRUE |
| **chr18:7854427:+** | INTRON | FALSE | FALSE | TRUE |
| **chr9:79169893:-** | INTRON | FALSE | FALSE | TRUE |
| **chrX:87724266:+** | INTRON | FALSE | FALSE | TRUE |
| **chr4:10936754:-** | INTERGENIC | FALSE | FALSE | TRUE |
| **chr6:140979694:-** | INTERGENIC | FALSE | FALSE | TRUE |
| **chr3:152322845:+** | INTRON | FALSE | FALSE | TRUE |
| **chr10:2822402:+** | INTERGENIC | FALSE | FALSE | TRUE |
| **chr2:29538824:+** | INTRON | FALSE | FALSE | TRUE |
| **chr1:80246735:+** | INTERGENIC | FALSE | FALSE | TRUE |
| **chr2:135475539:+** | INTRON | FALSE | FALSE | TRUE |
| **chr1:18339993:+** | INTRON | FALSE | FALSE | TRUE |
| **chr15:68294516:-** | UTR | FALSE | FALSE | TRUE |
| **chr17:64035391:-** | INTERGENIC | FALSE | FALSE | TRUE |
| **chr1:61495452:+** | INTRON | FALSE | FALSE | TRUE |
| **chr16:80887213:+** | INTRON | FALSE | FALSE | TRUE |
| **chr1:5444855:+** | INTERGENIC | FALSE | FALSE | TRUE |

| | | | | |
|---|---|---|---|---|
| **chr5:125103517:-** | INTRON | FALSE | FALSE | TRUE |
| **chr8:140632127:+** | INTRON | FALSE | FALSE | TRUE |
| **chrX:100642975:+** | INTERGENIC | FALSE | FALSE | TRUE |
| **chr9:107732814:-** | INTERGENIC | FALSE | FALSE | TRUE |
| **chr5:9106593:+** | INTRON | FALSE | FALSE | TRUE |
| **chr7:98585219:+** | INTERGENIC | FALSE | FALSE | TRUE |
| **chr8:67559005:-** | INTRON | FALSE | FALSE | TRUE |
| **chr8:135603147:+** | INTRON | FALSE | FALSE | TRUE |
| **chr8:136917323:+** | INTRON | FALSE | FALSE | TRUE |
| **chr2:107717804:+** | INTRON | FALSE | FALSE | TRUE |
| **chr8:3834030:+** | INTRON | FALSE | FALSE | TRUE |
| **chr8:124724411:-** | INTRON | FALSE | FALSE | TRUE |
| **chr2:230124521:-** | INTRON | FALSE | FALSE | TRUE |
| **chr4:138486819:+** | INTRON | FALSE | FALSE | TRUE |
| **chr5:127785864:-** | INTRON | FALSE | FALSE | TRUE |
| **chr13:93922985:+** | INTRON | FALSE | FALSE | TRUE |
| **chr3:170017499:-** | INTERGENIC | FALSE | FALSE | TRUE |
| **chr5:39896908:+** | INTRON | FALSE | FALSE | TRUE |
| **chr12:23708267:+** | INTRON | FALSE | FALSE | TRUE |
| **chr3:72899650:+** | INTRON | FALSE | FALSE | TRUE |
| **chr15:73640368:-** | INTERGENIC | FALSE | FALSE | TRUE |
| **chr10:92211469:+** | INTRON | FALSE | FALSE | TRUE |
| **chr5:75274185:-** | INTERGENIC | FALSE | FALSE | TRUE |
| **chr10:97481160:-** | INTRON | FALSE | FALSE | TRUE |
| **chr5:36281847:+** | INTRON | FALSE | FALSE | TRUE |
| **chr1:63453999:-** | INTRON | FALSE | FALSE | TRUE |
| **chr4:84081637:+** | INTRON | FALSE | FALSE | TRUE |
| **chr14:23718783:+** | INTRON | FALSE | FALSE | TRUE |
| **chr3:177012935:-** | INTERGENIC | FALSE | FALSE | TRUE |
| **chr11:41111773:-** | INTRON | FALSE | FALSE | TRUE |
| **chr22:48364199:+** | INTERGENIC | FALSE | FALSE | TRUE |
| **chr12:77850328:-** | INTRON | FALSE | FALSE | TRUE |
| **chr14:71719321:-** | INTRON | FALSE | FALSE | TRUE |
| **chr7:158760063:-** | CDS | FALSE | FALSE | TRUE |
| **chr11:25974947:+** | INTERGENIC | FALSE | FALSE | TRUE |
| **chr7:50418382:-** | INTERGENIC | FALSE | FALSE | TRUE |
| **chr11:78995305:-** | INTRON | FALSE | FALSE | TRUE |
| **chr8:106769342:-** | INTRON | FALSE | FALSE | TRUE |
| **chr8:39018097:+** | INTRON | FALSE | FALSE | TRUE |
| **chr22:12605583:-** | INTRON | FALSE | FALSE | TRUE |
| **chr7:112962944:+** | INTRON | FALSE | FALSE | TRUE |

| | | | | |
|---|---|---|---|---|
| **chr5:81968846:-** | INTERGENIC | FALSE | FALSE | TRUE |
| **chr12:16620181:+** | INTERGENIC | FALSE | FALSE | TRUE |
| **chr6:38702242:+** | INTRON | FALSE | FALSE | TRUE |
| **chr12:112226275:+** | INTRON | FALSE | FALSE | TRUE |
| **chr4:126285432:+** | INTERGENIC | FALSE | FALSE | TRUE |
| **chr1:150533133:-** | INTERGENIC | FALSE | FALSE | TRUE |
| **chr7:155128241:-** | INTERGENIC | FALSE | FALSE | TRUE |
| **chr15:38882470:-** | INTRON | FALSE | FALSE | TRUE |
| **chr7:17199011:+** | INTRON | FALSE | FALSE | TRUE |
| **chrX:133404135:+** | INTRON | FALSE | FALSE | TRUE |
| **chr13:40935984:-** | INTRON | FALSE | FALSE | TRUE |
| **chr5:149907552:+** | INTRON | FALSE | FALSE | TRUE |
| **chr2:96038453:+** | INTRON | FALSE | FALSE | TRUE |
| **chr2:95786128:-** | INTERGENIC | FALSE | FALSE | TRUE |
| **chr15:71184426:-** | INTRON | FALSE | FALSE | TRUE |
| **chr5:156458700:-** | INTRON | FALSE | FALSE | TRUE |
| **chr16:77305939:-** | INTRON | FALSE | FALSE | TRUE |
| **chr12:46665539:-** | INTRON | FALSE | FALSE | TRUE |
| **chr6:70146957:+** | INTRON | FALSE | FALSE | TRUE |
| **chr4:172164138:+** | INTRON | FALSE | FALSE | TRUE |
| **chr1:195798493:+** | INTERGENIC | FALSE | FALSE | TRUE |
| **chr15:39938981:-** | INTRON | FALSE | FALSE | TRUE |
| **chr3:174975723:-** | INTRON | FALSE | FALSE | TRUE |
| **chr13:75365021:-** | INTRON | FALSE | FALSE | TRUE |
| **chr10:116231942:-** | INTRON | FALSE | FALSE | TRUE |
| **chr8:136524991:+** | INTRON | FALSE | FALSE | TRUE |
| **chr3:159679560:-** | INTRON | FALSE | FALSE | TRUE |
| **chr1:15246676:+** | INTRON | FALSE | FALSE | TRUE |
| **chr17:78831994:+** | INTRON | FALSE | FALSE | TRUE |
| **chr10:131341624:+** | INTERGENIC | FALSE | FALSE | TRUE |
| **chr13:60002899:-** | INTRON | FALSE | FALSE | TRUE |
| **chr13:96277558:-** | INTRON | FALSE | FALSE | TRUE |
| **chr11:73138577:-** | INTRON | FALSE | FALSE | TRUE |
| **chr4:110999998:+** | INTRON | FALSE | FALSE | TRUE |
| **chr5:33525771:-** | UTR | FALSE | FALSE | TRUE |
| **chr2:42674155:+** | INTRON | FALSE | FALSE | TRUE |
| **chr20:62983446:+** | INTERGENIC | FALSE | FALSE | TRUE |
| **chr4:2335916:-** | INTRON | FALSE | FALSE | TRUE |
| **chr9:108863673:+** | UTR | FALSE | FALSE | TRUE |
| **chr13:109411523:+** | INTERGENIC | FALSE | FALSE | TRUE |
| **chr7:71865257:-** | INTRON | FALSE | FALSE | TRUE |

| | | | | |
|---|---|---|---|---|
| **chr11:59881043:-** | INTRON | FALSE | FALSE | TRUE |
| **chr7:72541690:+** | INTERGENIC | FALSE | FALSE | TRUE |
| **chr2:187069296:-** | INTRON | FALSE | FALSE | TRUE |
| **chr8:94770722:+** | INTRON | FALSE | FALSE | TRUE |
| **chr3:29554662:+** | INTRON | FALSE | FALSE | TRUE |
| **chr1:80112830:-** | INTERGENIC | FALSE | FALSE | TRUE |
| **chr11:32013036:+** | INTRON | FALSE | FALSE | TRUE |
| **chr6:73404747:-** | CDS | FALSE | FALSE | TRUE |
| **chr7:144043770:+** | INTERGENIC | FALSE | FALSE | TRUE |
| **chr22:21601628:+** | INTRON | FALSE | FALSE | TRUE |
| **chr1:203102519:-** | INTRON | FALSE | FALSE | TRUE |
| **chr6:4500417:+** | INTRON | FALSE | FALSE | TRUE |
| **chr3:179561722:+** | INTERGENIC | FALSE | FALSE | TRUE |
| **chr18:58700739:-** | INTRON | FALSE | FALSE | TRUE |
| **chr7:94508590:+** | INTERGENIC | FALSE | FALSE | TRUE |
| **chr12:92080426:-** | INTRON | FALSE | FALSE | TRUE |
| **chr3:60461946:-** | INTRON | FALSE | FALSE | TRUE |
| **chr10:54895164:-** | INTRON | FALSE | FALSE | TRUE |
| **chr8:7990730:+** | INTRON | FALSE | FALSE | TRUE |
| **chr8:7319632:-** | INTRON | FALSE | FALSE | TRUE |
| **chr3:29765542:-** | INTRON | FALSE | FALSE | TRUE |
| **chr16:16437155:-** | INTERGENIC | FALSE | FALSE | TRUE |
| **chr16:15319165:+** | INTERGENIC | FALSE | FALSE | TRUE |
| **chr16:14809621:-** | INTERGENIC | FALSE | FALSE | TRUE |
| **chr7:81831254:+** | INTERGENIC | FALSE | FALSE | TRUE |
| **chr3:74501330:+** | INTRON | FALSE | FALSE | TRUE |
| **chr2:97935061:-** | INTRON | FALSE | FALSE | TRUE |
| **chr9:128553626:-** | INTRON | FALSE | FALSE | TRUE |
| **chr21:25842893:-** | INTERGENIC | FALSE | FALSE | TRUE |
| **chr3:42168301:-** | INTRON | FALSE | FALSE | TRUE |
| **chr12:24008756:+** | INTRON | FALSE | FALSE | TRUE |
| **chr1:183074477:+** | INTRON | FALSE | FALSE | TRUE |
| **chr1:187350840:+** | INTRON | FALSE | FALSE | TRUE |
| **chr1:117536366:-** | INTERGENIC | FALSE | FALSE | TRUE |
| **chr3:18838418:+** | INTRON | FALSE | FALSE | TRUE |
| **chr7:40172360:-** | INTRON | FALSE | FALSE | TRUE |
| **chr13:42289736:-** | INTRON | FALSE | FALSE | TRUE |
| **chr21:36724825:-** | INTRON | FALSE | FALSE | TRUE |
| **chr3:141439486:-** | INTRON | FALSE | FALSE | TRUE |
| **chr1:180705754:+** | INTRON | FALSE | FALSE | TRUE |
| **chr4:97025519:-** | INTERGENIC | FALSE | FALSE | TRUE |

| | | | | |
|---|---|---|---|---|
| **chr6:116917430:+** | INTRON | FALSE | FALSE | TRUE |
| **chr17:79322484:-** | INTRON | FALSE | FALSE | TRUE |
| **chr8:100739890:-** | INTERGENIC | FALSE | FALSE | TRUE |
| **chr15:96562267:+** | INTERGENIC | FALSE | FALSE | TRUE |
| **chr2:176661714:-** | INTRON | FALSE | FALSE | TRUE |
| **chr5:50622491:+** | INTERGENIC | FALSE | FALSE | TRUE |
| **chr6:29869983:+** | INTRON | FALSE | FALSE | TRUE |
| **chr9:98904039:-** | INTERGENIC | FALSE | FALSE | TRUE |
| **chr5:63956318:-** | INTERGENIC | FALSE | FALSE | TRUE |
| **chr19:51264263:+** | INTRON | FALSE | FALSE | TRUE |
| **chr10:76577827:-** | INTERGENIC | FALSE | FALSE | TRUE |
| **chrX:141634419:-** | INTRON | FALSE | FALSE | TRUE |
| **chr10:79192559:-** | INTRON | FALSE | FALSE | TRUE |
| **chr6:23243827:+** | INTERGENIC | FALSE | FALSE | TRUE |
| **chr6:123026099:-** | INTRON | FALSE | FALSE | TRUE |
| **chr2:172013043:+** | INTRON | FALSE | FALSE | TRUE |
| **chr21:19672890:-** | INTERGENIC | FALSE | FALSE | TRUE |
| **chr1:226002495:+** | INTERGENIC | FALSE | FALSE | TRUE |
| **chrY:25046731:+** | INTRON | FALSE | FALSE | TRUE |
| **chrY:24623377:-** | INTRON | FALSE | FALSE | TRUE |
| **chrY:22989636:-** | INTRON | FALSE | FALSE | TRUE |
| **chr8:8716057:-** | INTRON | FALSE | FALSE | TRUE |
| **chr16:72089530:-** | INTRON | FALSE | FALSE | TRUE |
| **chr14:49297017:+** | INTERGENIC | FALSE | FALSE | TRUE |
| **chr8:103461430:-** | INTRON | FALSE | FALSE | TRUE |
| **chr3:157807738:+** | INTERGENIC | FALSE | FALSE | TRUE |
| **chr7:111080831:+** | INTRON | FALSE | FALSE | TRUE |
| **chr14:70619020:+** | INTRON | FALSE | FALSE | TRUE |
| **chr8:68916386:+** | INTRON | FALSE | FALSE | TRUE |
| **chr7:131490597:+** | UTR | FALSE | FALSE | TRUE |
| **chr10:114316101:+** | INTRON | FALSE | FALSE | TRUE |
| **chr2:10208509:+** | INTRON | FALSE | FALSE | TRUE |
| **chr19:21323920:-** | INTRON | FALSE | FALSE | TRUE |
| **chr2:29189197:-** | INTRON | FALSE | FALSE | TRUE |
| **chr3:42607219:+** | INTRON | FALSE | FALSE | TRUE |
| **chr4:72313110:+** | INTRON | FALSE | FALSE | TRUE |
| **chr2:76733958:-** | INTERGENIC | FALSE | FALSE | TRUE |
| **chr15:48663239:+** | INTERGENIC | FALSE | FALSE | TRUE |
| **chr13:106941512:+** | INTERGENIC | FALSE | FALSE | TRUE |
| **chr2:61791814:-** | INTERGENIC | FALSE | FALSE | TRUE |
| **chr3:162676203:-** | INTERGENIC | FALSE | FALSE | TRUE |

| | | | | |
|---|---|---|---|---|
| **chr12:123120404:-** | INTRON | FALSE | FALSE | TRUE |
| **chrY:14118466:+** | INTERGENIC | FALSE | FALSE | TRUE |
| **chr18:13939054:+** | INTERGENIC | FALSE | FALSE | TRUE |
| **chr8:9137968:+** | UTR | FALSE | FALSE | TRUE |
| **chr2:186972651:-** | INTERGENIC | FALSE | FALSE | TRUE |
| **chr15:35635251:+** | INTRON | FALSE | FALSE | TRUE |
| **chr5:112363975:+** | INTRON | FALSE | FALSE | TRUE |
| **chr9:85471789:-** | INTERGENIC | FALSE | FALSE | TRUE |
| **chr5:136511930:-** | INTRON | FALSE | FALSE | TRUE |
| **chr18:45171263:+** | INTRON | FALSE | FALSE | TRUE |
| **chr12:28438729:-** | INTRON | FALSE | FALSE | TRUE |
| **chr6:160922994:+** | INTRON | FALSE | FALSE | TRUE |
| **chr9:91964805:+** | INTERGENIC | FALSE | FALSE | TRUE |
| **chr2:14199826:-** | INTRON | FALSE | FALSE | TRUE |
| **chr12:27273308:+** | INTRON | FALSE | FALSE | TRUE |
| **chr11:71205848:-** | INTRON | FALSE | FALSE | TRUE |
| **chr8:113521296:+** | INTERGENIC | FALSE | FALSE | TRUE |
| **chr12:47144984:-** | INTRON | FALSE | FALSE | TRUE |
| **chrX:134254753:-** | INTERGENIC | FALSE | FALSE | TRUE |
| **chr20:25024099:-** | INTRON | FALSE | FALSE | TRUE |
| **chr22:19409225:-** | INTRON | FALSE | FALSE | TRUE |
| **chrX:10848005:-** | INTERGENIC | FALSE | FALSE | TRUE |
| **chr12:82740945:+** | INTRON | FALSE | FALSE | TRUE |
| **chr10:105173122:+** | INTRON | FALSE | FALSE | TRUE |
| **chr16:66239201:+** | INTERGENIC | FALSE | FALSE | TRUE |
| **chr11:82837694:+** | INTRON | FALSE | FALSE | TRUE |
| **chr17:79532679:+** | INTRON | FALSE | FALSE | TRUE |
| **chr11:113655328:+** | INTERGENIC | FALSE | FALSE | TRUE |
| **chr2:2406505:-** | INTERGENIC | FALSE | FALSE | TRUE |
| **chr6:164302282:-** | INTERGENIC | FALSE | FALSE | TRUE |
| **chr4:118748570:+** | INTRON | FALSE | FALSE | TRUE |
| **chr13:69761749:-** | INTRON | FALSE | FALSE | TRUE |
| **chr9:108542377:+** | INTERGENIC | FALSE | FALSE | TRUE |
| **chr21:44803291:+** | INTRON | FALSE | FALSE | TRUE |
| **chr18:29061135:-** | INTERGENIC | FALSE | FALSE | TRUE |
| **chr2:99544614:-** | INTERGENIC | FALSE | FALSE | TRUE |
| **chr8:17422368:+** | INTERGENIC | FALSE | FALSE | TRUE |
| **chr8:86614145:+** | INTRON | FALSE | FALSE | TRUE |
| **chr18:33797629:-** | INTERGENIC | FALSE | FALSE | TRUE |
| **chrX:110803775:+** | INTERGENIC | FALSE | FALSE | TRUE |
| **chr7:47706114:+** | INTERGENIC | FALSE | FALSE | TRUE |

| | | | | |
|---|---|---|---|---|
| **chrX:125403138:-** | INTERGENIC | FALSE | FALSE | TRUE |
| **chr6:137373569:+** | INTERGENIC | FALSE | FALSE | TRUE |
| **chr9:2442584:+** | INTRON | FALSE | FALSE | TRUE |
| **chr2:38440727:-** | INTRON | FALSE | FALSE | TRUE |
| **chr4:120862641:-** | INTRON | FALSE | FALSE | TRUE |
| **chr3:71708412:-** | INTRON | FALSE | FALSE | TRUE |
| **chr18:49179486:+** | INTRON | FALSE | FALSE | TRUE |
| **chr4:67850557:-** | INTRON | FALSE | FALSE | TRUE |
| **chr6:152014881:+** | INTRON | FALSE | FALSE | TRUE |
| **chr17:69726542:+** | INTRON | FALSE | FALSE | TRUE |
| **chr4:91213232:+** | INTRON | FALSE | FALSE | TRUE |
| **chr18:2788316:-** | INTRON | FALSE | FALSE | TRUE |
| **chr14:30394498:-** | INTERGENIC | FALSE | FALSE | TRUE |
| **chr4:61091170:+** | INTERGENIC | FALSE | FALSE | TRUE |
| **chr5:19486166:+** | INTRON | FALSE | FALSE | TRUE |
| **chrX:50492374:-** | INTERGENIC | FALSE | FALSE | TRUE |
| **chr8:15984304:+** | INTRON | FALSE | FALSE | TRUE |
| **chr10:108245317:-** | INTERGENIC | FALSE | FALSE | TRUE |
| **chr9:98446972:+** | INTRON | FALSE | FALSE | TRUE |
| **chr4:67655122:+** | INTRON | FALSE | FALSE | TRUE |
| **chr17:65117589:-** | INTRON | FALSE | FALSE | TRUE |
| **chr10:25229834:+** | INTRON | FALSE | FALSE | TRUE |
| **chr7:105442912:+** | INTRON | FALSE | FALSE | TRUE |
| **chr19:40671622:+** | INTRON | FALSE | FALSE | TRUE |
| **chr5:151185659:-** | INTRON | FALSE | FALSE | TRUE |
| **chr6:144969052:-** | INTERGENIC | FALSE | FALSE | TRUE |
| **chr20:10477084:-** | INTRON | FALSE | FALSE | TRUE |
| **chr5:104710437:-** | INTRON | FALSE | FALSE | TRUE |
| **chr2:111072354:+** | INTRON | FALSE | FALSE | TRUE |
| **chr1:27878521:-** | INTRON | FALSE | FALSE | TRUE |
| **chr7:78588029:+** | INTRON | FALSE | FALSE | TRUE |
| **chr4:138093440:-** | INTRON | FALSE | FALSE | TRUE |
| **chr10:132482199:-** | INTERGENIC | FALSE | FALSE | TRUE |
| **chr7:7883344:-** | INTRON | FALSE | FALSE | TRUE |
| **chr8:57911192:-** | INTRON | FALSE | FALSE | TRUE |
| **chr15:94998203:-** | INTRON | FALSE | FALSE | TRUE |
| **chr17:56928428:+** | INTERGENIC | FALSE | FALSE | TRUE |
| **chr3:108843746:+** | INTRON | FALSE | FALSE | TRUE |
| **chr2:50348219:+** | INTRON | FALSE | FALSE | TRUE |
| **chr7:76656000:+** | INTERGENIC | FALSE | FALSE | TRUE |
| **chr1:83190819:+** | INTERGENIC | FALSE | FALSE | TRUE |

| | | | | |
|---|---|---|---|---|
| **chr3:51847737:-** | INTERGENIC | FALSE | FALSE | TRUE |
| **chr6:1213064:+** | INTRON | FALSE | FALSE | TRUE |
| **chr1:147958672:-** | INTRON | FALSE | FALSE | TRUE |
| **chr21:19101074:-** | INTERGENIC | FALSE | FALSE | TRUE |
| **chr9:21397800:-** | INTRON | FALSE | FALSE | TRUE |
| **chr15:70296249:-** | INTERGENIC | FALSE | FALSE | TRUE |
| **chr4:61585026:-** | INTRON | FALSE | FALSE | TRUE |
| **chrX:85415966:-** | INTERGENIC | FALSE | FALSE | TRUE |
| **chr20:62032097:-** | INTRON | FALSE | FALSE | TRUE |
| **chr17:14941355:+** | INTRON | FALSE | FALSE | TRUE |
| **chr11:27239748:+** | INTERGENIC | FALSE | FALSE | TRUE |
| **chr10:101196043:+** | INTERGENIC | FALSE | FALSE | TRUE |
| **chr10:2885603:-** | INTERGENIC | FALSE | FALSE | TRUE |
| **chr2:72195397:-** | INTRON | FALSE | FALSE | TRUE |
| **chr8:49883330:-** | INTERGENIC | FALSE | FALSE | TRUE |
| **chr7:125056107:+** | INTRON | FALSE | FALSE | TRUE |
| **chr14:20671773:-** | INTERGENIC | FALSE | FALSE | TRUE |
| **chr1:220491690:+** | INTERGENIC | FALSE | FALSE | TRUE |
| **chr10:101056305:+** | INTERGENIC | FALSE | FALSE | TRUE |
| **chr20:52936004:-** | INTERGENIC | FALSE | FALSE | TRUE |
| **chr5:6166446:-** | INTERGENIC | FALSE | FALSE | TRUE |
| **chr16:72063081:+** | INTRON | FALSE | FALSE | TRUE |
| **chr16:72054478:+** | INTRON | FALSE | FALSE | TRUE |
| **chr8:34088787:+** | INTERGENIC | FALSE | FALSE | TRUE |
| **chr1:50069452:+** | INTRON | FALSE | FALSE | TRUE |
| **chr12:91522747:+** | INTRON | FALSE | FALSE | TRUE |
| **chr4:31237895:+** | INTERGENIC | FALSE | FALSE | TRUE |
| **chr8:47574819:+** | INTRON | FALSE | FALSE | TRUE |
| **chr13:91402323:+** | INTRON | FALSE | FALSE | TRUE |
| **chr15:76947725:-** | INTRON | FALSE | FALSE | TRUE |
| **chr18:79893316:+** | INTRON | FALSE | FALSE | TRUE |
| **chr6:114178921:-** | INTRON | FALSE | FALSE | TRUE |
| **chr3:52828267:-** | INTRON | FALSE | FALSE | TRUE |
| **chr11:128440479:-** | INTERGENIC | FALSE | FALSE | TRUE |
| **chr13:78094502:+** | INTRON | FALSE | FALSE | TRUE |
| **chr15:101828486:-** | INTERGENIC | FALSE | FALSE | TRUE |
| **chr20:59871424:-** | INTRON | FALSE | FALSE | TRUE |
| **chr2:13557903:-** | INTRON | FALSE | FALSE | TRUE |
| **chr3:96093793:-** | INTERGENIC | FALSE | FALSE | TRUE |
| **chr3:48120363:-** | INTERGENIC | FALSE | FALSE | TRUE |
| **chr7:105379305:+** | INTRON | FALSE | FALSE | TRUE |

| | | | | |
|---|---|---|---|---|
| **chr8:118403028:-** | INTRON | FALSE | FALSE | TRUE |
| **chr4:112459652:-** | INTERGENIC | FALSE | FALSE | TRUE |
| **chr3:124398649:+** | INTRON | FALSE | FALSE | TRUE |
| **chr11:125507751:+** | INTRON | FALSE | FALSE | TRUE |
| **chr4:1461320:+** | INTERGENIC | FALSE | FALSE | TRUE |
| **chr2:197860987:-** | INTRON | FALSE | FALSE | TRUE |
| **chr11:8963751:+** | INTRON | FALSE | FALSE | TRUE |
| **chr2:85949254:-** | INTERGENIC | FALSE | FALSE | TRUE |
| **chr2:124575251:+** | INTRON | FALSE | FALSE | TRUE |
| **chr13:32169851:-** | INTRON | FALSE | FALSE | TRUE |
| **chr11:29673020:+** | INTRON | FALSE | FALSE | TRUE |
| **chr8:42435522:-** | INTRON | FALSE | FALSE | TRUE |
| **chrX:151004528:-** | INTERGENIC | FALSE | FALSE | TRUE |
| **chr16:74660522:-** | INTRON | FALSE | FALSE | TRUE |
| **chr10:84904721:+** | INTERGENIC | FALSE | FALSE | TRUE |
| **chr7:92124406:-** | INTRON | FALSE | FALSE | TRUE |
| **chr5:62647507:+** | INTERGENIC | FALSE | FALSE | TRUE |
| **chr1:40702011:-** | INTRON | FALSE | FALSE | TRUE |
| **chr3:177362372:+** | INTERGENIC | FALSE | FALSE | TRUE |
| **chr11:99970785:-** | INTRON | FALSE | FALSE | TRUE |
| **chr20:4577008:+** | INTERGENIC | FALSE | FALSE | TRUE |
| **chr7:148727992:-** | INTRON | FALSE | FALSE | TRUE |
| **chr11:120829249:-** | INTRON | FALSE | FALSE | TRUE |
| **chr9:74451678:-** | INTERGENIC | FALSE | FALSE | TRUE |
| **chr1:241695343:-** | INTRON | FALSE | FALSE | TRUE |
| **chr2:3562563:+** | INTERGENIC | FALSE | FALSE | TRUE |
| **chr12:67234563:+** | INTERGENIC | FALSE | FALSE | TRUE |
| **chr20:19402770:+** | INTRON | FALSE | FALSE | TRUE |
| **chr5:105083581:-** | INTRON | FALSE | FALSE | TRUE |
| **chr21:17898384:-** | INTERGENIC | FALSE | FALSE | TRUE |
| **chr2:65614219:-** | INTRON | FALSE | FALSE | TRUE |
| **chr8:83087050:+** | INTERGENIC | FALSE | FALSE | TRUE |
| **chr20:25600722:-** | INTERGENIC | FALSE | FALSE | TRUE |
| **chr17:61019543:-** | INTRON | FALSE | FALSE | TRUE |
| **chr7:1994178:+** | INTRON | FALSE | FALSE | TRUE |
| **chr1:20849498:-** | CDS | FALSE | FALSE | TRUE |
| **chr7:108187461:-** | INTRON | FALSE | FALSE | TRUE |
| **chr14:82315832:+** | INTERGENIC | FALSE | FALSE | TRUE |
| **chr5:60641716:+** | INTRON | FALSE | FALSE | TRUE |
| **chr11:63670287:+** | INTRON | FALSE | FALSE | TRUE |
| **chrX:87601293:+** | INTRON | FALSE | FALSE | TRUE |

| | | | | |
|---|---|---|---|---|
| **chr13:42818205:+** | INTERGENIC | FALSE | FALSE | TRUE |
| **chr3:143560312:-** | INTRON | FALSE | FALSE | TRUE |
| **chr9:124116663:-** | INTERGENIC | FALSE | FALSE | TRUE |
| **chr3:191825946:-** | INTERGENIC | FALSE | FALSE | TRUE |
| **chr2:85321608:+** | UTR | FALSE | FALSE | TRUE |
| **chr6:169393629:+** | INTERGENIC | FALSE | FALSE | TRUE |
| **chrX:125940582:+** | INTERGENIC | FALSE | FALSE | TRUE |
| **chr8:93941481:+** | INTRON | FALSE | FALSE | TRUE |
| **chr5:97808288:+** | INTRON | FALSE | FALSE | TRUE |
| **chr4:154953781:+** | INTERGENIC | FALSE | FALSE | TRUE |
| **chr2:172535162:-** | INTRON | FALSE | FALSE | TRUE |
| **chrX:88605657:-** | INTERGENIC | FALSE | FALSE | TRUE |
| **chr2:73902994:-** | UTR | FALSE | FALSE | TRUE |
| **chr7:78676947:+** | INTRON | FALSE | FALSE | TRUE |
| **chr4:68899301:+** | INTERGENIC | FALSE | FALSE | TRUE |
| **chr12:51247585:-** | INTRON | FALSE | FALSE | TRUE |
| **chr8:75692508:-** | INTERGENIC | FALSE | FALSE | TRUE |
| **chr8:132122180:+** | UTR | FALSE | FALSE | TRUE |
| **chr15:69821110:+** | INTRON | FALSE | FALSE | TRUE |
| **chr12:81145791:-** | INTRON | FALSE | FALSE | TRUE |
| **chr10:61992198:-** | INTRON | FALSE | FALSE | TRUE |
| **chr22:40864413:+** | INTRON | FALSE | FALSE | TRUE |
| **chr10:108894124:-** | INTERGENIC | FALSE | FALSE | TRUE |
| **chr12:26230472:+** | INTRON | FALSE | FALSE | TRUE |
| **chr20:43183117:-** | INTRON | FALSE | FALSE | TRUE |
| **chr4:25961166:+** | INTERGENIC | FALSE | FALSE | TRUE |
| **chr4:94405477:+** | INTERGENIC | FALSE | FALSE | TRUE |
| **chr5:12995157:-** | INTRON | FALSE | FALSE | TRUE |
| **chr10:12720974:+** | INTRON | FALSE | FALSE | TRUE |
| **chr20:51588441:-** | INTERGENIC | FALSE | FALSE | TRUE |
| **chr20:21432667:-** | INTERGENIC | FALSE | FALSE | TRUE |
| **chr5:133563803:-** | INTRON | FALSE | FALSE | TRUE |
| **chr3:162042463:+** | INTERGENIC | FALSE | FALSE | TRUE |
| **chr5:20315519:+** | INTRON | FALSE | FALSE | TRUE |
| **chr18:65963554:-** | INTERGENIC | FALSE | FALSE | TRUE |
| **chr11:84828064:+** | INTRON | FALSE | FALSE | TRUE |
| **chr8:125343254:+** | INTRON | FALSE | FALSE | TRUE |

**Table S4: Summary of insertion events detected at the CCR5 on-target site by Bionano *De Novo* Assembly.** The expected size of HDR-mediated FIX transgene insertion is 1948 bp. For each sample analyzed by OGM, the size and Variant Allele Fraction (VAF) for insertion events at the CCR5 on-target site as identified by Bionano *De Novo* Assembly are listed. The insertion fraction was independently determined by ddPCR to provide an orthogonal measurement.

<table>
<tr><th>Sample</th><th>Expected insertion size (bp)</th><th>Observed insertion size (bp)</th><th>VAF</th><th>ddPCR insertion fraction</th></tr>
<tr><td>Donor 1</td><td rowspan="3">1948</td><td>1955</td><td>0.49</td><td>0.49</td></tr>
<tr><td>Donor 2</td><td>1935</td><td>0.40</td><td>0.46</td></tr>
<tr><td>Donor 3</td><td>1966</td><td>0.43</td><td>0.45</td></tr>
</table>

**Table S5: Summary of insertion events detected at the CCR5 on-target site by Bionano Rare Variant Analysis.** For each sample analyzed by OGM, sizes and molecule counts for large insertion events at the CCR5 on-target site as identified by Bionano Rare Variant Analysis are listed.

| Sample | Observed insertion size (bp) | Molecule count |
|---|---|---|
| Donor 1 | 5484 | 25 |
| | 8799 | 5 |
| | 12884 | 5 |
| Donor 2 | 5534 | 18 |
| | 8866 | 9 |
| Donor 3 | 5470 | 21 |
| | 8740 | 9 |

**Table S6: *F9* Transgene FIX Production from BE-101 Cells from Scale Down Process with and without the RNP Complex**

| Donor ID | Process Day | Process | Scale | hFIX (ng/mL) | % FIX (no RNP/RNP) |
|---|---|---|---|---|---|
| 23031 | Day 11 | BE-101 | Scale Down | 353 | 1.9 |
| | | BE-101 no RNP | | 7 | |
| | Day 14 | BE-101 | | 676 | 3.3 |
| | | BE-101 no RNP | | 22 | |
| 23037 | Day 11 | BE-101 | | 382 | 1.8 |
| | | BE-101 no RNP | | 7 | |
| | Day 14 | BE-101 | | 1182 | 1.9 |
| | | BE-101 no RNP | | 22 | |
| 23154 | Day 14 | BE-101 no RNP | Clinical Scale | 9 | <1% of average at scale |

**Table S7: Hematology data at Day 28 in BE-101 GLP toxicology study in NOG-hIL6 mice**

| Test | Units | Vehicle Average (n=5) | BE-101 Donor 1 Average (n=3) | BE-101 Donor 2 Average (n=3) |
|---|---|---|---|---|
| Leukocyte count | $x10^3$ /µL | 1.1 | 0.9 | 0.7 |
| Red blood cell count | $X10^6$/µL | 8.7 | 8.4 | 8.9 |
| Hemoglobin | g /dL | 13.3 | 13.1 | 13.7 |
| Hematocrit | % | 48.2 | 46.5 | 49.9 |
| Mean cell volume | fL | 55.4 | 55.5 | 56.1 |
| Mean corpuscular hemoglobin | pg | 15.3 | 15.5 | 15.4 |
| Mean corpuscular hemoglobin concentration | g / dL | 27.7 | 28.0 | 27.4 |
| Red blood cell distribution width | % | 15.2 | 15.5 | 15.0 |
| Platelet count | $x10^3$ /µL | 1703.2 | 1601.3 | 1788.0 |
| Mean platelet volume | fL | 6.4 | 6.5 | 5.9 |
| % neutrophils | % | 70.6 | 70.0 | 74.0 |
| % lymphocytes | % | 21.6 | 25.3 | 22.0 |
| % monocytes | % | 5.0 | 2.7 | 2.7 |
| % eosinophils | % | 2.4 | 2.0 | 1.3 |
| % basophils | % | 0.4 | 0.7 | 0.0 |
| % band cells | % | 0.0 | 0.0 | 0.0 |
| Neutrophil count | $x10^3$ /µL | 0.8 | 0.6 | 0.5 |
| Lymphocyte count | $x10^3$ /µL | 0.2 | 0.2 | 0.2 |
| Monocyte count | $x10^3$ /µL | 0.1 | 0.0 | 0.0 |
| Eosinophil count | $x10^3$ /µL | 0.0 | 0.0 | 0.0 |
| Basophil count | $x10^3$ /µL | 0.0 | 0.0 | 0.0 |
| Band cell count | $x10^3$ /µL | 0.0 | 0.0 | 0.0 |
| % reticulocytes | % | 4.5 | 5.2 | 3.7 |
| Reticulocyte count | $x10^9$/L | 391.4 | 433.9 | 325.2 |

**Table S8: Coagulation data at Day 28 in BE-101 GLP toxicology study in NOG-hIL6 mice**

| Test | Units | Vehicle Average (n=5) | BE-101 Donor 1 Average (n=3) | BE-101 Donor 2 Average (n=3) |
|---|---|---|---|---|
| Prothrombin time | Seconds | 12.7 | 12.9 | 13.3 |
| Activated partial thromboplastin time | Seconds | 25.4 | 27.1 | 28 |
| Fibrinogen | mg/dL | 180.4 | 184.3 | 215.7 |

**Table S9: Clinical chemistry data at Day 28 in BE-101 GLP toxicology study in NOG-hIL6 mice**

| Test | Units | Vehicle Average (n=5) | BE-101 Donor 1 Average (n=3) | BE-101 Donor 2 Average (n=3) |
|---|---|---|---|---|
| Albumin | g/dL | 2.8 | 2.8 | 2.8 |
| Total protein | g/dL | 4.7 | 4.5 | 4.6 |
| Alkaline phosphatase | U/L | 47.2 | 40.3 | 43.8 |
| Alanine transaminase | U/L | 67.8 | 64.0 | 65.9 |
| Aspartate aminotransferase | U/L | 136.2 | 265.7 | 200.9 |
| Creatine kinase | U/L | 291.8 | 319.5 | 305.7 |
| Total bilirubin | mg/dL | 0.2 | 0.3 | 0.3 |
| Direct bilirubin | mg/dL | 0.1 | 0.1 | 0.1 |
| Blood urea nitrogen | mg/dL | 24.4 | 26.7 | 25.5 |
| Creatinine* | mg/dL | 0.2 | 0.2 | 0.2 |
| Calcium | mg/dL | 9.8 | 9.5 | 9.6 |
| Cholesterol | mg/dL | 76.2 | 66.7 | 71.4 |
| Glucose | mg/dL | 139.2 | 149.3 | 144.3 |
| Phosphorus | mg/dL | 7.7 | 7.5 | 7.6 |
| Gamma-glutamyl transferase | U/L | 0.0 | 0.0 | 0.0 |
| Total carbon dioxide | mEq/L | 29.0 | 38.0 | 33.5 |
| Sodium | mEq/L | 152.2 | 150.7 | 151.4 |
| Potassium | mEq/L | 4.8 | 6.0 | 5.4 |
| Chloride | mEq/L | 113.8 | 114.0 | 113.9 |
| Triglycerides | mg/dL | 92.8 | 88.7 | 90.7 |
| Globulin | g/dL | 1.9 | 1.8 | 1.8 |
| Albumin/Globulin | Ratio | 1.5 | 1.6 | 1.6 |
| Blood Urea Nitrogen / Creatinine | Ratio | 140.0 | 142.5 | 141.3 |
| Indirect bilirubin | mg/dL | 0.1 | 0.2 | 0.2 |
| Anion Gap | mEq/L | 14.2 | 5.0 | 9.6 |

*all creatinine samples were 0.2 or less, with some reported as “less than 0.2” due to the limits of the assay

**Table S10: Tissues examined microscopically in 28-day GLP toxicology study**

| | | | |
|---|---|---|---|
| Spleen | Liver | Kidney | Lung |
| Heart (with aorta) | Brain | Spinal column | Adrenal glands |
| Testes | Lymph nodes | Stomach | Intestine |
| Pancreas | Esophagus | Trachea | Skin |
| Skeletal muscle | Eyes | Gallbladder | Salivary gland |
| Thyroids | Tongue | Urinary bladder | Pituitary gland |
| Bone (femur) | | | |

**Table S11: Hematology data at 28 weeks: non-GLP pharmacology study in NOG-hIL6 mice**

| Test | Units | Vehicle Group Average (n=6) | BE-101 Group Average (n=4) |
|---|---|---|---|
| Leukocyte count | K/µL | 1.42 | 1.25 |
| Red blood cell count | M/µL | 8.80 | 9.22 |
| Hemoglobin | g/dL | 13.18 | 13.53 |
| Hematocrit | % | 43.48 | 45.48 |
| Mean cell volume | fL | 49.67 | 49.50 |
| Red blood cell distribution width | % | 20.38 | 20.58 |
| Mean corpuscular hemoglobin | pg | 15.02 | 14.65 |
| Mean corpuscular hemoglobin concentration | g/dL | 30.32 | 29.75 |
| Platelet count | K/µL | 955.50 | 1540.25 |
| Mean platelet volume | fL | 6.90 | 6.63 |
| % neutrophils | % | 58.67 | 60.40 |
| % lymphocytes | % | 31.80 | 26.40 |
| % monocytes | % | 9.02 | 12.28 |
| % eosinophils | % | 0.52 | 0.78 |
| % basophils | % | 0.00 | 0.15 |
| Neutrophil count | Absolute number /µL | 869.33 | 773.75 |
| Lymphocyte count | Absolute number /µL | 416.83 | 303.25 |
| Monocyte count | Absolute number /µL | 123.83 | 159.50 |
| Eosinophil count | Absolute number /µL | 6.67 | 11.00 |
| Basophil count | Absolute number /µL | 0.00 | 2.75 |
| % reticulocytes | % | 4.12 | 5.03 |
| Reticulocyte count | K/µL | 363.50 | 462.75 |

**Table S12: Clinical chemistry data at 28 weeks: non-GLP pharmacology study in NOG-hIL6 mice**

| Test | units | Vehicle Group Average (n=4) | BE-101 Group Average (n=4) |
|---|---|---|---|
| Albumin | g/dL | 3.00 | 2.95 |
| Total protein | g/dL | 4.83 | 4.63 |
| Alkaline phosphatase | U/L | 54.00 | 44.00 |
| Aspartate aminotransferase | U/L | 170.50 | 73.25 |
| Alanine transaminase | U/L | 90.75 | 34.50 |
| Creatine kinase | U/L | 536.00 | 171.50 |
| Total bilirubin | mg/dL | 0.10 | 0.10 |
| Direct bilirubin | mg/dL | 0.00 | 0.00 |
| Globulin | g/dL | 1.83 | 1.68 |
| Blood urea nitrogen | mg/dL | 23.25 | 22.00 |
| Creatinine | mg/dL | 0.08 | 0.08 |
| Calcium | mg/dL | 9.30 | 9.50 |
| Cholesterol | mg/dL | 44.00 | 44.25 |
| Glucose | mg/dL | 134.50 | 149.50 |
| Phosphorus | mg/dL | 6.65 | 5.93 |
| Total carbon dioxide | mmol/L | 19.25 | 20.25 |
| Sodium | mmol/L | 147.33 | 148.50 |
| Chloride | mmol/L | 108.33 | 109.50 |
| Potassium | mmol/L | 4.33 | 5.00 |
| Albumin/Globulin | Ratio | 1.68 | 1.75 |
| Triglycerides | mg/dL | | |
| Blood Urea Nitrogen / Creatinine | Ratio | 88.75 | 80.00 |
| Indirect bilirubin | mg/dL | 0.10 | 0.10 |

**Table S13: Tissues examined microscopically in long-term pharmacology studies**

| Tissue Examined | Number of BE-101 treated Animals |
|---|---|
| Spleen | 15 |
| Heart | 15 |
| Lung | 15 |
| Liver | 15 |
| Gallbladder | 6 |
| Kidney | 15 |
| Brain | 15 |
| Spinal column | 15 |
| Esophagus | 6 |
| Aorta | 6 |
| Bone (femur) | 6 |
| Muscle | 6 |
| Stomach | 6 |
| Pancreas | 6 |
| Large and small intestine | 6 |
| Thyroid | 6 |
| Bladder | 6 |
| Uterus | 6 |

**Table S14: Guide ribonucleic acid (gRNA) sequence**

| **Description** | **Sequence** |
|---|---|
| gCCR5_232 | 5’-CAATGTGTCAACTCTTGACA-3' |

**Table S15: Human *F9* gene (hyperactive R338L, Padua variant and Malmö T148A variant) expression cassette sequencing**

| Description | Nucleotide Sequence |
|---|---|
| *F9 Padua* | 5’ -<br>GCCACCATGCAGAGGGTGAACATGATCATGGCTGAGAGCCCTGGCCTGAT<br>CACCATCTGCCTGCTGGGCTACCTGCTGTCTGCTGAGTGCACTGTGTTCCT<br>GGACCATGAGAATGCCAACAAGATCCTGAACAGGCCCAAGAGATACAAC<br>TCTGGCAAGCTGGAGGAGTTTGTGCAGGGCAACCTGGAGAGGGAGTGCA<br>TGGAGGAGAAGTGCAGCTTTGAGGAGGCCAGGGAGGTGTTTGAGAACAC<br>TGAGAGGACCACTGAGTTCTGGAAGCAGTATGTGGATGGGGACCAGTGT<br>GAGAGCAACCCCTGCCTGAATGGGGGCAGCTGCAAGGATGACATCAACA<br>GCTATGAGTGCTGGTGCCCCTTTGGCTTTGAGGGCAAGAACTGTGAGCTG<br>GATGTGACCTGCAACATCAAGAATGGCAGATGTGAGCAGTTCTGCAAGA<br>ACTCTGCTGACAACAAGGTGGTGTGCAGCTGCACTGAGGGCTACAGGCTG<br>GCTGAGAACCAGAAGAGCTGTGAGCCTGCTGTGCCATTCCCATGTGGCAG<br>AGTGTCTGTGAGCCAGACCAGCAAGCTGACCAGGGCTGAGGCTGTGTTCC<br>CTGATGTGGACTATGTGAACAGCACTGAGGCTGAAACCATCCTGGACAAC<br>ATCACCCAGAGCACCCAGAGCTTCAATGACTTCACCAGGGTGGTGGGGG<br>GGGAGGATGCCAAGCCTGGCCAGTTCCCCTGGCAAGTGGTGCTGAATGGC<br>AAGGTGGATGCCTTCTGTGGGGGCAGCATTGTGAATGAGAAGTGGATTGT<br>GACTGCTGCCCACTGTGTGGAGACTGGGGTGAAGATCACTGTGGTGGCTG<br>GGGAGCACAACATTGAGGAGACTGAGCACACTGAGCAGAAGAGGAATGT<br>GATCAGGATCATCCCCCACCACAACTACAATGCTGCCATCAACAAGTACA<br>ACCATGACATTGCCCTGCTGGAGCTGGATGAGCCCCTGGTGCTGAACAGC<br>TATGTGACCCCCATCTGCATTGCTGACAAGGAGTACACCAACATCTTCCT<br>GAAGTTTGGCTCTGGCTATGTGTCTGGCTGGGGCAGGGTGTTCCACAAGG<br>GCAGGTCTGCCCTGGTGCTGCAGTACCTGAGGGTGCCCCTGGTGGACAGG<br>GCCACCTGCCTGCTGAGCACCAAGTTCACCATCTACAACAACATGTTCTG<br>TGCTGGCTTCCATGAGGGGGGCAGGGACAGCTGCCAGGGGGACTCTGGG<br>GGCCCCCATGTGACTGAGGTGGAGGGCACCAGCTTCCTGACTGGCATCAT<br>CAGCTGGGGGGAGGAGTGTGCCATGAAGGGCAAGTATGGCATCTACACC<br>AAAGTCTCCAGATATGTGAACTGGATCAAGGAGAAGACCAAGCTGACCT<br>GA- 3’ |

**Table S16: Primers used for Inference of CRISPR Edits (ICE)**

| Description | Nucleotide Sequence |
|---|---|
| **Primer 1 Sequence (5’-3’)** | 5’ -GCAGCAAACCTTCCCTTCACTAC- 3’ |
| **Primer 2 Sequence (5’-3’)** | 5’ -AGGATTCCCGAGTAGCAGATGAC- 3’ |
| **Sequencing Primer Sequence (5’-3’)** | 5’-GGGTGGAACAAGATGGATTATC- 3’ |

**Table S17: Primers used for ddPCR analysis of targeted integration of the *F9* transgene**

| Primer | Sequence | Target | Assay |
|---|---|---|---|
| TJM_221 | 5' -CATCGCATTGTCTGAGTAGG- 3' | BGH polyA | Integrated transgene |
| TJM_349 | 5' -CAGTGGATCGGGTGTAAAC- 3' | CCR5 locus downstream of homology arm | |
| TJM_346 | 5' -TCGGGAGCCTCTTGCTGGAAAATAGAA- 3' | CCR5 locus downstream of homology arm (FAM probe) | |
| TJM_376 | 5' -CCACATCAGAAGGAAGACTAC- 3' | C-C chemokine receptor-like 2 (CCRL2) locus | Reference gene |
| TJM_230 | 5' -GCTGTATGAATCCAGGTCC- 3' | CCRL2 locus | |
| TJM_307 | 5' -TGTTTCCTCCAGGATAAGGCAGCTGT- 3' | CCRL2 locus (HEX probe) | |

**Supplemental Figure**

**Figure S1: Structural variant identification by Bionano *De Novo* Assembly.** Circos plots depicting structural variants (SVs) as identified by Bionano *De Novo* Assembly. in three lots of BE-101 (top) with magnification of the on-target insertion site on chromosome 3 (below). The outermost numerical track corresponds to the chromosome number. The cytoband information is shown in the black-and-white banding pattern with red lines indicating the centromere. The next circle indicates SVs that were uniquely identified in engineered cells but not donor-matched unengineered controls. Insertion events are represented by green dots. The next circle shows copy number variations. The inner space indicates translocation events found uniquely in engineered cells. Insertion events are summarized in Table S5.

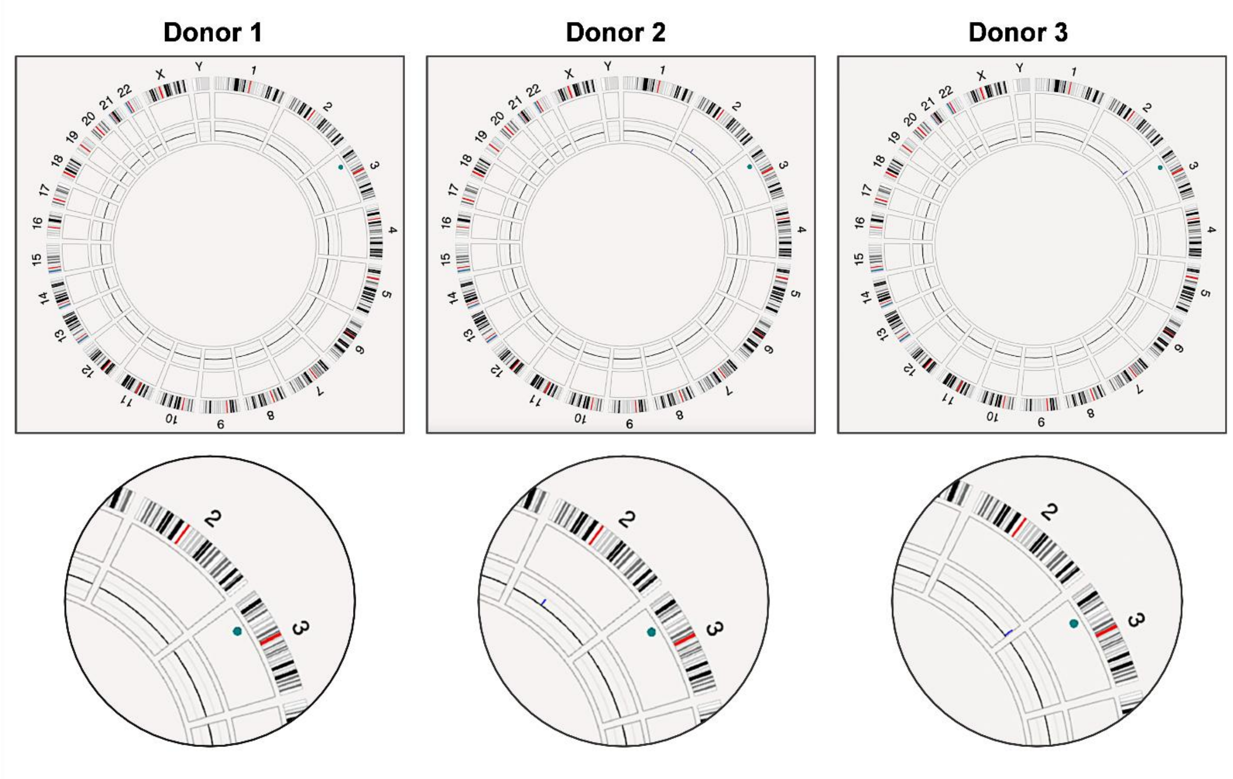

**Figure S2. Secretion of human IgM after a single IV administration of BE-101 to NOG-hIL6 mice.** Each data point represents the mean at that timepoint. The error bars indicate the Standard Error of the Mean (SEM). The lower limit of quantitation (LLOQ) of human IgM is 1.22 ng/mL.

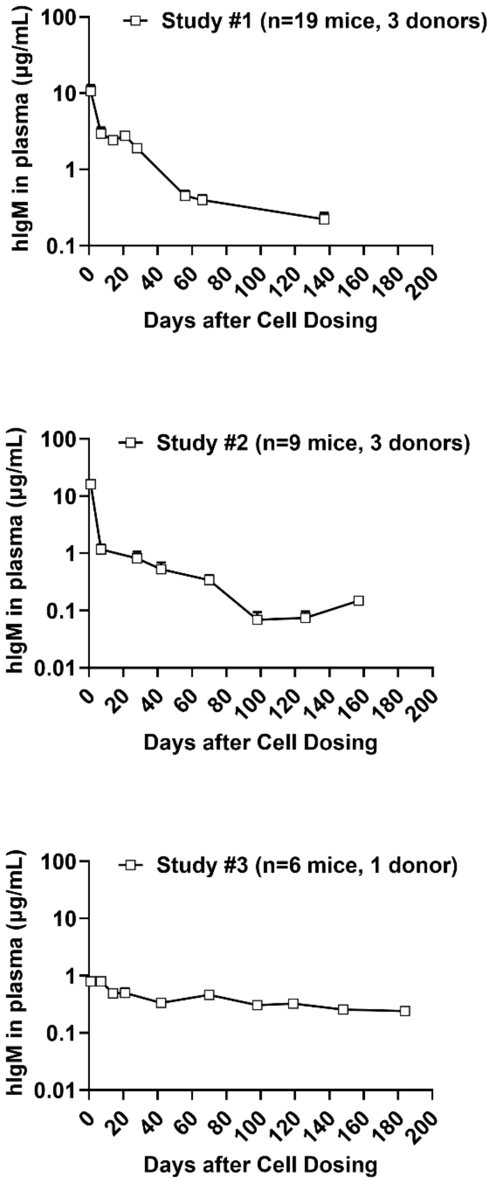

**Figure S3. Human IgM protein in plasma after repeated intravenous administration of BE-101 to NOG-hIL6 mice.** A) NOG-hIL6 mice (n =4) were dosed with 20e6 total viable cells of BE-101 on day 0 and redosed on day 49 of study. IgG and IgM levels were measured in blood plasma after 1 day post BE-101 administration and then weekly after each dose. Closed circles represent blood plasma measurements after single dose and open circles represent levels after redose. Each data point represents the mean at that timepoint. The error bars indicate the Standard Error of the Mean (SEM). The lower limit of quantitation (LLOQ) of human IgM is 0.195 ng/mL.

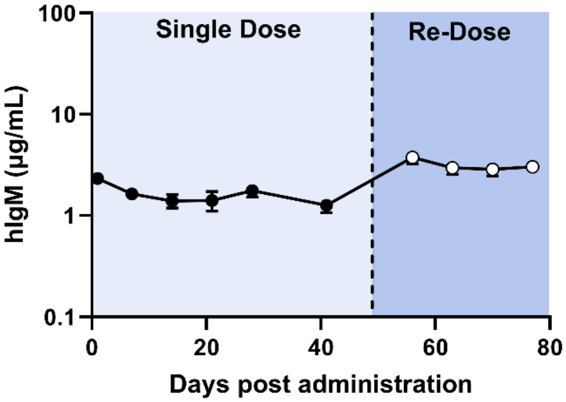

### Figure S4. Average body weight over time in 28 day GLP toxicity study

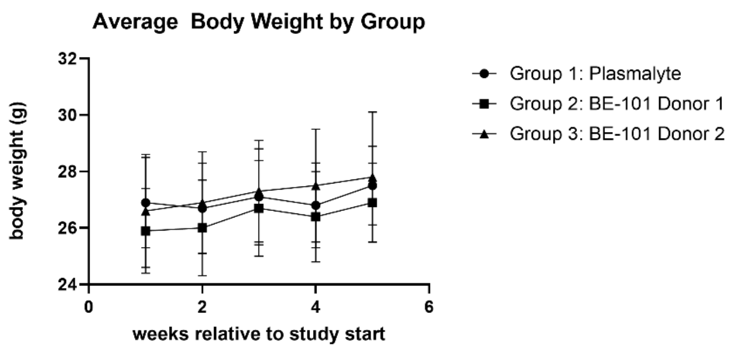

Body weights were collected 1 week prior to dosing and each week until termination

**Figure S5. Average body weight over time: 28-week non-GLP pharmacology study**

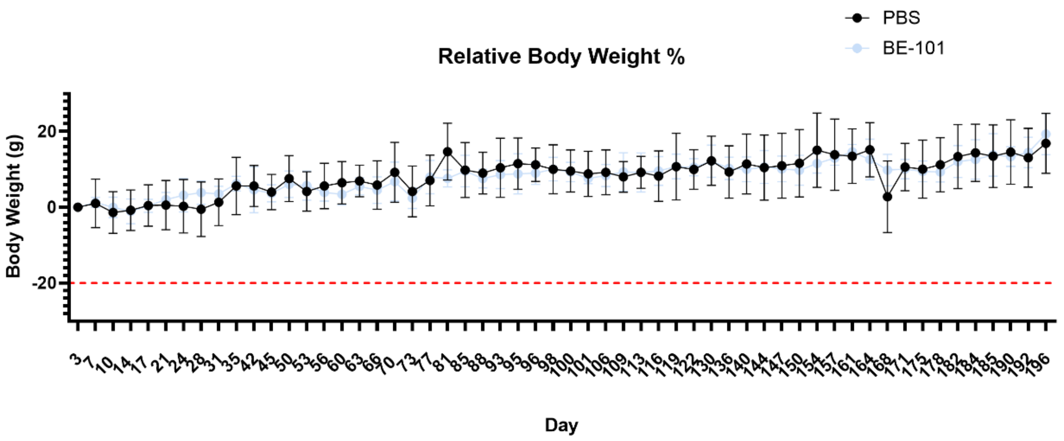
